# Spike-frequency dependent inhibition and excitation of neural activity by high-frequency ultrasound

**DOI:** 10.1101/2020.06.01.128710

**Authors:** Martin Loynaz Prieto, Kamyar Firouzi, Butrus T. Khuri-Yakub, Daniel V. Madison, Merritt Maduke

**Affiliations:** Department of Molecular and Cellular Physiology, Stanford University; E. L. Ginzton Laboratory, Stanford University

## Abstract

Ultrasound can modulate action-potential firing *in vivo* and *in vitro*, but the mechanistic basis of this phenomenon is not well understood. To address this problem, we used patch-clamp recording to quantify the effects of focused, high-frequency (43 MHz) ultrasound on evoked action potential firing in CA1 pyramidal neurons in acute rodent hippocampal brain slices. We find that ultrasound can either inhibit or potentiate firing in a spike-frequency-dependent manner: at low (near-threshold) input currents and low firing frequencies, ultrasound inhibits firing, while at higher input currents and higher firing frequencies, ultrasound potentiates firing. The net result of these two competing effects is that ultrasound increases the threshold current for action potential firing, the slope of frequency-input curves, and the maximum firing frequency. In addition, ultrasound slightly hyperpolarizes the resting membrane potential, decreases action potential width, and increases the depth of the afterhyperpolarization. All of these results can be explained by the hypothesis that ultrasound activates a sustained potassium conductance. According to this hypothesis, increased outward potassium currents hyperpolarize the resting membrane potential and inhibit firing at near-threshold input currents, but potentiate firing in response to higher input currents by limiting inactivation of voltage-dependent sodium channels during the action potential. This latter effect is a consequence of faster action-potential repolarization, which limits inactivation of voltage-dependent sodium channels, and deeper (more negative) afterhyperpolarization, which increases the rate of recovery from inactivation. Based on these results we propose that ultrasound activates thermosensitive and mechanosensitive two-pore-domain potassium (K2P) channels, through heating or mechanical effects of acoustic radiation force. Finite-element modelling of the effects of ultrasound on brain tissue suggests that the effects of ultrasound on firing frequency are caused by a small (less than 2°C) increase in temperature, with possible additional contributions from mechanical effects

**SUMMARY:** Prieto et al. describe how ultrasound can either inhibit or potentiate action potential firing in hippocampal pyramidal neurons and demonstrate that these effects can be explained by increased potassium conductance.

## INTRODUCTION

Ultrasound can non-invasively modulate action potential activity in neurons *in vivo* and *in vitro*, with improved depth penetration and spatial resolution relative to other non-invasive neuromodulation modalities, and it may therefore become an important new technology in basic and clinical neuroscience (Fry et al., 1958; Gavrilov et al., 1996; Tufail et al., 2010; Bystritsky et al., 2011; Fomenko et al., 2018; Blackmore et al., 2019). Investigation of this phenomenon has predominantly focused on low-frequency ultrasound (defined here as less than 3 MHz, although there is no firmly defined boundary between “high” and “low” frequency in the neuromodulation field), but higher ultrasound frequencies have also been shown to modulate action potential firing *in vitro* (Menz et al., 2013; Menz et al., 2019) and to directly modulate ion channel activity in heterologous systems (Kubanek et al., 2016; Prieto et al., 2018). The focus on lower frequency ultrasound is understandable, since envisioned clinical applications involving transcranial focused ultrasound have been a primary motivation for research on ultrasound neuromodulation, and loss of ultrasound power due to attenuation in the skull limits these applications to low frequency ultrasound. For applications in which transmission through the skull does not impose limits on frequency, such as *in vitro* studies, neuromodulation in the peripheral nervous system (Downs et al., 2018; Cotero et al., 2019; Zachs et al., 2019), neuromodulation using subcranial implants, or neuromodulation in experimental animal model systems involving craniotomies or acoustically transparent cranial windows, high frequencies have a distinct advantage in terms of the greater spatial resolution that can be achieved. Even for *in vivo* applications in human subjects, however, the spatial resolutions that can be achieved with low-frequency, transcranial ultrasound neuromodulation are on the order of millimeters, making ultrasound neuromodulation superior in this respect to other, more established forms of non-invasive brain stimulation (Tyler et al., 2018).

These applications motivate investigation of the fundamental physical, cellular, and molecular mechanisms underlying neuromodulation, which are all not well understood for either high- or low-frequency ultrasound. It remains an open question to what extent these mechanisms overlap for high- and low-frequency ultrasound neuromodulation. In terms of the basic physical mechanism by which acoustic energy is transduced into effects on biological tissue, most proposed mechanisms for ultrasound neuromodulation involve either heating due to absorption of acoustic energy (Hand, 1998), mechanical effects of acoustic radiation force (Duck, 1998; Sarvazyan et al., 2010), or effects of cavitation (the nucleation, growth, oscillation, and sometimes collapse, of microscopic gas bubbles) (Leighton, 1998; Wu and Nyborg, 2008; Krasovitski et al., 2011; Plaksin et al., 2016). Of these, the first two increase with acoustic frequency, while the probability of cavitation decreases with acoustic frequency. There are also many unanswered questions at the cellular level. Both excitatory and inhibitory effects of ultrasound have been observed using direct or indirect measures of neural activity at the population level (Bystritsky et al., 2011; Blackmore et al., 2019), but it is unclear whether the direct effect of ultrasound at the cellular level is excitatory or inhibitory. Of course, the answer to this question could depend on any number of possible relevant biological and experimental variables, such as species, tissue, specific neural subtype, ultrasound stimulus parameters, or whether effects on intrinsic or evoked activity are measured. For example, a cellular-level excitatory effect, specific to GABAergic interneurons, could produce an inhibitory effect at the population level. This leads to the question of whether certain subpopulations of neurons are more sensitive to ultrasound than others, and if so, what are the specific molecular mechanisms underlying the differences in sensitivity. Do certain ion channels respond directly to ultrasound? What biophysical properties might account for the sensitivity of these channels to ultrasound, and how might cell-type specific differences in the density and localization of these channels, and the way in which they interact with other ion channels to regulate action potential firing, produce differences in the response to ultrasound?

One reason there are so many outstanding questions regarding ultrasound neuromodulation is that patch-clamp recordings of the effects of ultrasound on action-potential firing in neurons have been unavailable. At low ultrasound frequencies, we (Prieto et al., 2018) and others (Tyler et al., 2008) have found that patch-clamp seals are extremely unstable in the presence of ultrasound at low frequencies, precluding detailed, mechanistic studies of ultrasound neuromodulation with this technique, which provides quantitative information on action-potential timing and dynamics unobtainable with other techniques. However, we have previously shown that stable patch-clamp recordings can be achieved using ultrasound at the frequency of 43 MHz (Prieto et al., 2018). Here, we use patch-clamp recording to measure the effects of ultrasound at 43 MHz and 50 W/cm^2^ on action-potential firing in response to injected current in pyramidal neurons of the CA1 layer of the hippocampus in acute rodent brain slices. We find that ultrasound has a bidirectional, spike-frequency dependent effect on excitability, and that this and other observed neurophysiological effects of ultrasound can be explained by activation by ultrasound of a steady K^+^ current, resembling that of two pore domain potassium channels (K2P channels).

## MATERIALS AND METHODS

### Slice preparation

Brain slices were prepared from male Sprague-Dawley rats, 35-50 days old. Rats were anesthetized with isofluorane and decapitated, and the brain was immediately removed and placed in ice-cold artificial cerebral spinal fluid (ACSF), bubbled with 95% O_2_ and 5% CO_2_. The hippocampus was dissected out and placed on the slicing apparatus, consisting of a manual micrometer and a gravity-driven vertical slicing mechanism, with the CA1 layer oriented approximately parallel to the slicing blade. Slices (500 microns thick) were prepared and then stored in a humidified chamber with an atmosphere of 95% O_2_/5% CO_2_, resting on a square of filter paper placed on a dish of ACSF. Slices were used within 1-6 hours of slice preparation. Animals were handled in accordance with protocols approved by Stanford University’s Institutional Animal Care and Use Committee.

### Ultrasound

Continuous wave ultrasound at 43 MHz and 50 W/cm^2^ was applied to brain slices using a set-up similar to that we previously used for our experiments on cultured cells (Prieto et al., 2018), except that the tissue was visualized with a dissecting microscope at low magnification. The bottom of the experimental chamber was a 25-micron film of polystyrene, plasma-treated with a Harrick plasma cleaner (Harrick Plasma, Ithaca, NY). Ultrasound was transmitted from below (through the polystyrene film) with the sound beam perpendicular to the bottom of the chamber. The 43 MHz transducer was a custom-built device, calibrated as described previously (Prieto et al., 2013), excited using an ENI 403LA (37 dB) amplifier (ENI, Rochester, NY). The focal volume of the transducer is approximately a cylinder 90 microns in diameter by 500 microns long, and the focal distance is approximately 4.2 mm. The set-up was based on the stage from a Zeiss Axioskop-2 microscope (Zeiss Microscopes, Jena, Germany), with the housing for the sub-stage condenser modified to accommodate the transducer, such that the position of the transducer could be adjusted using the controls for alignment of the condenser, and the position of the tissue sample relative to the transducer could be adjusted with the microscope stage. The transducer was coupled to the polystyrene film at the bottom of the experimental chamber using a small volume of distilled water held in place by a rubber O-ring attached to the tip of the transducer with silicone grease. The focal volume of the transducer was aligned along the z-axis using a pulse-echo protocol, adjusting the height of the transducer to maximize the echo signal from the bottom of the empty chamber. The focus was aligned in the x-y plane by adding to the chamber a small volume of ACSF, barely sufficient to cover the bottom of the chamber, such that a thin layer of solution was spread over the bottom of the chamber. Ultrasound pulses, one second in length, were then applied, raising a mound of fluid at the focus of the transducer (due to the radiation force produced by reflection of the acoustic wave at the interface between the solution and the air above it (Duck, 1998)). The mound of fluid was then aligned in the x-y plane to the center of a reticle in one eyepiece of the dissecting microscope, and, after adding additional ACSF and the tissue sample to the chamber, the center of the reticle was aligned with the region of the tissue targeted for patch-clamp recording. The ultrasound intensity (50 W/cm^2^) is the spatial peak, pulse average intensity for the free field. The interval between ultrasound applications was at least 12 seconds.

### Electrophysiology

Current clamp recordings were performed using an Axon Instruments Axoclamp-2B amplifier operating in “Bridge” mode and Digidata 1330A digitizer with pClamp software (Molecular Devices, Sunnyvale, CA), except for the preliminary experiments in Supplemental Figure 1, which were performed with an Axon Instruments Axopatch 200B amplifier (Molecular Devices, Sunnyvale, CA). Patch-clamp recording was performed using a “blind-patch” approach (Blanton et al., 1989; Malinow and Tsien, 1990; Castaneda-Castellanos et al., 2006), in which the recording pipette was positioned above the CA1 layer of the hippocampus, as identified visually at low magnification, and then slowly lowered into the tissue while applying positive pressure to the pipette and monitoring the pipette tip resistance in voltage-clamp mode. In the blind-patch approach, a small decrease in tip resistance is used to indicate possible contact of the pipette tip with a neuron in the absence of the usual visual cues. Typically, the first two instances of possible cell contact were not used, to avoid patching on cells at the surface of the tissue that may have been damaged during the slicing procedure. Gigaseals and the subsequent whole-cell recording configuration were obtained following the standard procedure in voltage-clamp mode before switching to current-clamp mode. In most experiments, slices were held in place with a Warner Instruments RC-22 slice anchor (Harvard Bioscience, Hamden, CT); the experiments in Supplemental Figure 1 and some of the experiments in Figure 1E were performed without a slice anchor. No obvious effects of the slice anchor on the ultrasound response were noted. Series resistance, monitored and compensated throughout the recording, was between 30 and 100 MΩ. All of the neurons used for experiments could be unambiguously identified as pyramidal cells by their distinct adaptive firing patterns in response to 2-s current steps. Current records were low-pass filtered at 10 kHz and sampled at 100 kHz. Brain slices were continuously perfused with ACSF (in mM: 119 NaCl, 2.5 KCl, 1.3 MgSO_4_, 2.5 CaCl_2,_ 1 NaH_2_PO_4_, 26.2 NaHCO_3_, 11 glucose), bubbled with 95% O_2_/5% CO_2_, at ∼100-250 mL/hour. The internal solution was (in mM): 126 K-gluconate, 4 KCl, 10 HEPES, 4 Mg-ATP, 0.3 Na_2_GTP, 10 Na-phosphocreatine, 10 sucrose, and 50 U/mL creatine phosphokinase (porcine), pH 7.2 (KOH). This internal solution contains an ATP-regenerating system (phosphocreatine and creatine phosphokinase) because we found that the strength of the response to ultrasound was unstable, gradually declining over the course of a recording unless the ATP-regenerating system was included (Supplemental Figure 2). Na-phosphocreatine was obtained from Abcam and creatine phosphokinase was obtained from EMD Millipore. All other salts and chemicals were obtained from either Sigma-Aldrich or Fisher Scientific. (For some of the preliminary recordings shown in Figure 1E, the creatine phosphokinase was omitted from the internal solution, or a different internal solution, containing 120 K-gluconate, 40 HEPES, 5 MgCl_2_, 0.3 Na_2_GTP, 2 Na_2_ATP, pH 7.2 (KOH) was used. Creatine phosphokinase was also omitted for the experiments in Supplemental Figure 4). Other than the reduction of the ultrasound response over time in the absence of creatine phosphokinase, no obvious differences in recordings with different internal solutions were noted.) Because creatine phosphokinase increases the viscosity of the solution, making it difficult to obtain gigaseals, a small volume of internal solution without the enzyme was added to the tip of the pipette (enough to fill approximately the first 3 mm of the tip) before back-filling the pipette with the enzyme-containing solution. Pipettes were pulled from thick walled glass and had resistances between 5 and 10 MΩ. Recordings were performed at room temperature (21-23°C), except for the experiments in Supplemental Figure 4, which were performed at near physiological temperature (30°C). For these experiments, the temperature of the external solution was regulated and monitored with a Warner Instruments CL-100 bipolar temperature controller equipped with a SC-20 in-line heater/cooler and a thermistor (Warner Instruments, Hamden, CT). The external solution was heated to 35-37°C while being bubbled with 95% O_2_/5% CO_2_ and then passed through the heater/cooler and cooled to achieve the target temperature of 30°C in the bath solution. (The external solution was cooled rather than heated to avoid loss of oxygen tension and formation of gas bubbles due to heating of oxygen-saturated solution.)

**Figure 1.**
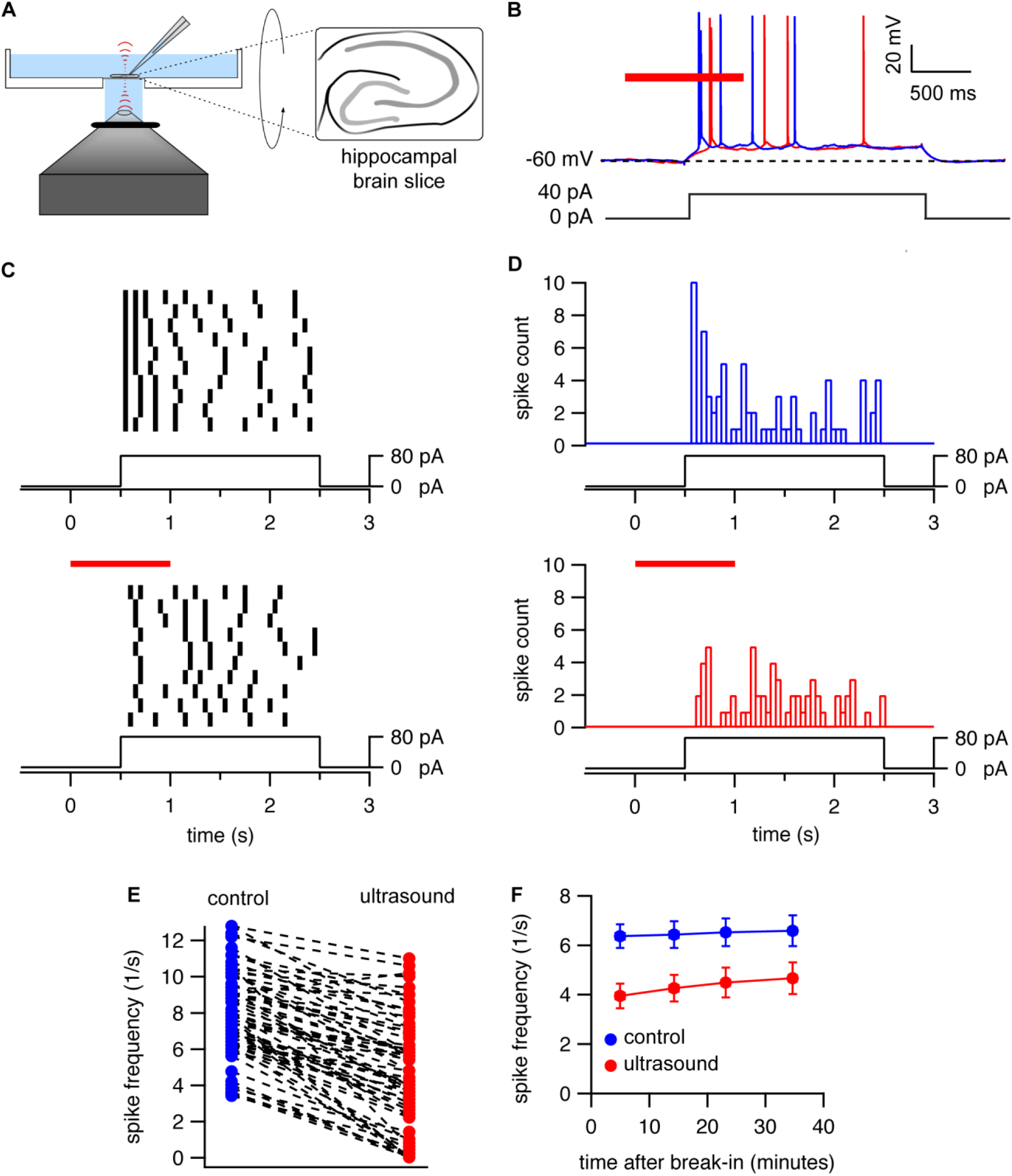
Consistent inhibition of action potential firing by high-frequency ultrasound. **A.** Diagram of experimental set-up. Ultrasound is applied to 500-micron hippocampal brain slices resting on a 25-micron film of polystyrene. The 43-MHz focused transducer is located below the experimental chamber, with ultrasound propagating perpendicular to the bottom of the recording chamber. **B.** Experimental protocol and example voltage traces showing inhibition of action potential firing by ultrasound. A 1-s, continuous-wave ultrasound pulse at 43 MHz and 50 W/cm^2^ (*red bar*) is applied 500 ms before the start of a 2-s current injection. Voltage traces are shown in the presence (*red*) and absence (*blue*) of the ultrasound stimulus. The *dashed line* indicates the resting membrane voltage. **C.** Example raster plots showing a consistent effect of ultrasound on firing frequency. The results of twenty consecutive trials of the protocol in panel B, alternating between the control (*top*) and ultrasound (*bottom*) conditions, are shown. The voltage traces were divided into 50-ms bins; a solid black bin indicates that an action potential occurred within that particular time bin. Time is relative to the start of the ultrasound pulse. **D.** Spike-time histograms prepared by summing the ten trials for the control (*top, blue bars*) and ultrasound (*bottom, red bars*) conditions from panel C. **E.** Summary of the effects of ultrasound for N = 66 cells. The average firing frequency during the first 500 ms of the current step is shown for the control (*blue*) and ultrasound (*red*) conditions. **F.** Stability of the ultrasound response. Mean (±SEM, N = 10 cells) spike frequencies during the first 500 ms of a current step in the presence (*red circles*) and absence (*blue circles*) of ultrasound for the protocol shown in panel B, as a function of time relative to break-in (establishment of whole-cell recording configuration). Spike frequencies were measured at various time points between 0 and 10, 10 and 20, 20 and 30, and 30 and 40 minutes after break-in. The x-values represent the mean start time for the protocol to measure spike frequencies (which comprised 2 minutes of recording time). The amplitude of the current step was adjusted over time to maintain spiking behavior as close as possible to that at the start of the experiment.

### Data analysis

Current records were analyzed in Igor Pro (Wavemetrics, Lake Oswego, OR) with user-written procedures. Action-potential threshold was defined as the point at which the first derivative of the voltage reached 4% of its peak value during the rising phase of the action potential. This quantitative criterion was previously found to correspond with action-potential thresholds as identified visually (Khaliq and Bean, 2010; Yamada-Hanff and Bean, 2015), and we found that it also works well with our data, using phase plots to visually confirm the threshold value. Action-potential height was defined as the difference between the action-potential peak and the action-potential threshold voltage. Action-potential width was measured at 50% of action potential height defined in this manner. Threshold current levels for action-potential firing were estimated based on a series of current steps in 10-pA increments. Frequency-input plots and action-potential parameters (height, width, latency, and interspike intervals) were determined from the average values of at least 3 trials each for the control and ultrasound conditions. Frequency-input trials were performed alternatingly for the control and ultrasound conditions, with the first condition tested varying randomly on a cell-by-cell basis. Average traces for analysis of the effects of ultrasound on membrane resting potential and membrane capacitance were derived from at least 3 voltage traces. Statistical significance was assessed using paired or unpaired two-tailed Student’s t-tests, with P < 0.05 defined as significant. Statistical analysis was performed in Microsoft Excel.

### Finite-element simulations

Finite-element were performed in COMSOL (COMSOL Inc., Palo Alto, CA, USA). The simulation domain had radially symmetric geometry and was 6 mm in the axial direction. The simulation domain contained four layers of different materials: a lower layer of water (4.2 mm thick in the axial direction), followed by a layer of polystyrene (25 microns thick), followed by a layer of brain tissue (500 microns thick), followed by an upper layer of water (1.275 mm thick). The width of the simulation domain in the axial direction was 1 mm (for simulation for acoustic pressure and heating) or 5 mm (for simulation of mechanical deformation) in the axial direction. A 940-micron diameter by 100-micron height arc on the lower axial boundary of the simulation domain represented the quartz lens of the transducer.

Simulations of acoustic pressure, heating, and static displacement in response to radiaton force were performed as described previously (Prieto et al., 2018). For simulation of dynamic tissue displacement in response to radiation force, the brain slice was modeled as a incompressible, linear viscoelastic material (Calhoun et al., 2019), characterized by Young’s modulus, Poisson’s ratio, and shear viscosity, loaded by the fluid layer above it. The polystyrene was modeled as a linear elastic material, because we determined in a series of simulations that including viscosity of the polystyrene had no effect on the tissue displacement. A time step of 0.1 ms was used for simulation of the dynamic tissue displacement. Material properties used for water, polystyrene, and brain tissue used in the simulation and sources for these values are given in Table 1. Additional details on mesh size, boundary conditions, and solver configurations are available in Prieto et al. (2018).

**Table 1.**
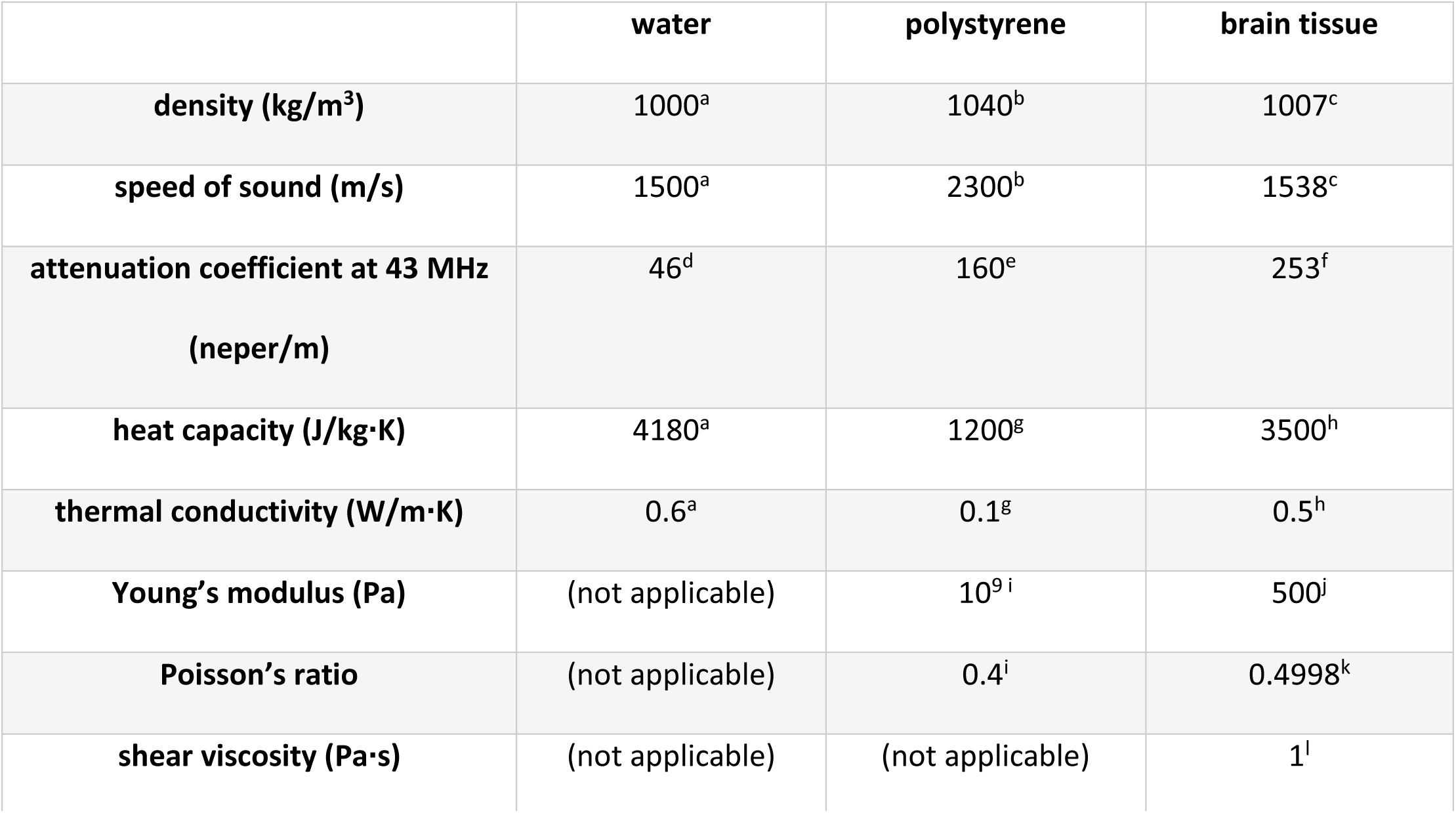
**Values of Material Properties Used in Finite-Element Simulations.** Sources for material properties are as follows: ^a^standard value; ^b^based on typical acoustic properties of plastics (Selfridge, 1985); ^c^following Menz et al. (2019), based on (Thijssen et al., 1985); ^d^ (Company, 1965); ^e^measured (Prieto et al., 2018); ^f^following Menz et al. (2019), based on (de Korte et al., 1994); ^g^based on typical thermal properties of plastics (Gaur and Wunderlich, 1982; Harper, 2006); ^h^typical values for soft tissues (Hand, 1998); ^i^based on typical mechanical properties of plastics (Harper, 2006); ^j^Menz et al. (2019), from measurements of ultrasound-induced displacement in the retina; ^k^tissue assumed to be incompressible for small deformations; ^l^see text.

## SUMMARY OF SUPPLEMENTAL MATERIAL

Supplemental materials include four figures showing: the effect of ultrasound at different intensities on action potential firing frequency (Supplemental Figure 1); stabilization of the response to ultrasound by an ATP-regenerating system in the internal solution (Supplemental Figure 2); effect of ultrasound on action potential height (Supplemental Figure 3); and the effects of ultrasound on action potential firing and waveform at near physiological temperature (30°C) (Supplemental Figure 4).

## RESULTS

We measured the effects of ultrasound on action-potential firing in hippocampal CA1 pyramidal cells using the set-up shown in Figure 1A (described in *Materials and Methods*). Throughout the experiments reported here, ultrasound was applied at 43 MHz and 50 W/cm^2^ as a 1-s, continuous-wave pulse. In an initial exploration of the effects of intensity on the ultrasound response, we determined that 50 W/cm^2^ had a sufficiently robust effect on firing frequency to permit quantitative investigation of this effect (Supplemental Figure 1), but we did not perform a detailed investigation of the intensity dependence. We chose to use continuous-wave ultrasound (without additional low-frequency modulation within the pulse) because continuous-wave ultrasound was previously found to be optimal for ultrasound neuromodulation of retinal ganglion cells at 43 MHz (Menz et al., 2013). With these ultrasound parameters, we found robust, reproducible inhibition of action-potential firing by ultrasound using the protocol illustrated in Figure 1B. In these experiments, a current-injection amplitude sufficient to induce firing at an average frequency of ∼4-12 Hz during the first 500 ms of the current step (corresponding to the overlap between the ultrasound stimulus and current step) was used. This range of firing frequencies is physiologically relevant and sufficient to detect either inhibition or potentiation of firing. With these experimental conditions, we established that the response to ultrasound is highly reproducible, both on a trial-by-trial basis within the same cell (Figure 1C-D), and between cells, with similar effects seen in over fifty cells (Figure 1E). In addition, the response to ultrasound was stable over the course of recordings lasting over 30 minutes (Figure 1F, Supplemental Figure 2); in a few cases where the patch seal lasted for over 90 minutes, the ultrasound response remained stable.

### Effects of ultrasound on frequency-input curves

To explore the effects of ultrasound on excitability over a wider range of firing frequencies, we generated frequency-input (f-i) curves comparing average firing frequencies as a function of input current in the presence and absence of ultrasound. An example f-i curve generated with the protocol illustrated in Figure 1B is shown in Figure 2A, along with example voltage traces in Figure 2B. The average spike frequency during the first 500 ms of the current step was compared to the spike frequency in the same time window in the absence of ultrasound. To compare the effects of ultrasound across neurons, we converted the f-i curves into plots of the relative increase or decrease in firing frequency as a function of the input current (Figure 2C-D). These data reveal two distinct regimes with contrasting inhibitory and excitatory ultrasound effects. At relatively low input currents, near the threshold for action-potential firing under this current stimulation protocol, ultrasound decreases the average firing frequency; while at relatively high input currents, well above the action-potential threshold, ultrasound increases the average firing frequency. Between these two regimes, there is a transitional region where there is little or no effect on average firing frequency, presumably due to the balance between competing inhibitory and excitatory effects. Other notable effects of ultrasound on the f-i curves are an increase in the threshold current for action-potential firing, an increase in the slope of the f-i relationship in the approximately linear region of the f-i curve, and an increase in the maximum firing frequency in the sublinear “plateau” region of the curve (Figure 2A). The mean (± SEM) slope of linear region of the f-i curve increased from 0.108 ± 0.007 Hz/pA in the control condition to 0.145 ± 0.012 Hz/pA in the ultrasound condition; and the mean maximum firing frequency increased from 23 ± 1 Hz in the control condition to 30 ± 2 Hz/pA in the ultrasound condition (N = 9).

**Figure 2.**
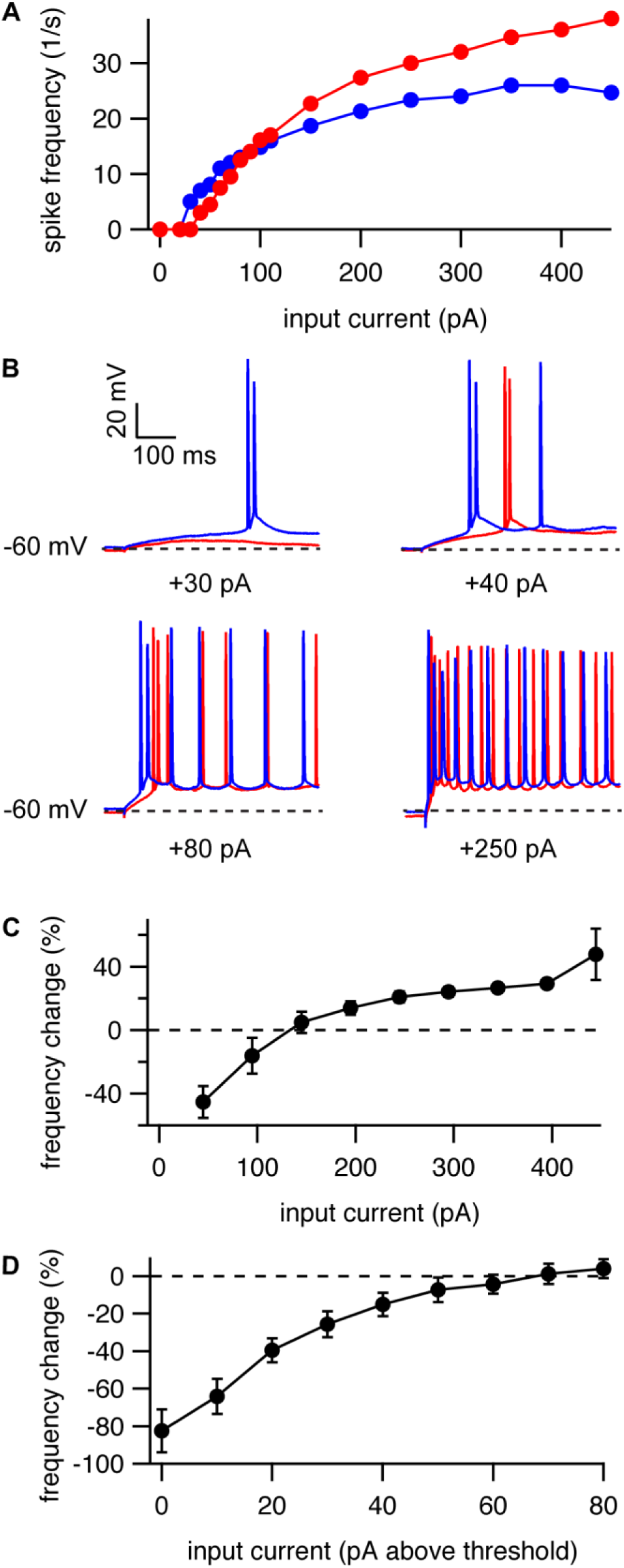
Ultrasound can inhibit or potentiate action-potential firing. **A.** Example frequency-input curve showing average firing frequency during the first 500 ms of a current step as a function of input current, with (*red*) and without (*blue*) a 1-s ultrasound pulse starting before 500 ms before the start of the current step. Each point represents the average of three trials on the same cell. **B.** Example voltage traces for the cell in A showing action potential firing during the first 500 ms of current steps to either +30, +40, +80, or +250 pA, with (*red*) and without (*blue*) ultrasound. The *dashed lines* indicate the approximate resting membrane voltage of −60 mV. **C.** Mean (±SEM, N = 9 cells) change in spike frequency in response to ultrasound as a function of input current. **D.** Inhibition is strongest near threshold. Mean (±SEM, N = 7 cells) change in spike frequency in response to ultrasound as a function of input current relative to the threshold current for action potential firing, for near-threshold currents.

Action-potential firing behavior is determined by the interaction between numerous K^+^, Na^+^, and other ionic currents (Madison and Nicoll, 1984; Bean, 2007). Some of these currents are clearly identified with specific ion channel subtypes, while the molecular identity of others is still uncertain. Thus, f-i curves are a complicated function of the density, localization, conductance, and kinetic properties of these channels/currents. Some currents inactivate relatively rapidly and only influence firing frequency during the initial response to a sustained depolarizing current step, while others show slow, voltage-dependent activation, and only influence firing frequency late in a current step; still other currents can influence firing frequency throughout a sustained depolarization. To explore the molecular basis of the response to ultrasound, we therefore generated a second set of f-i curves with ultrasound applied 1 s after the start of a 3-s current step (Figure 3A).

**Figure 3.**
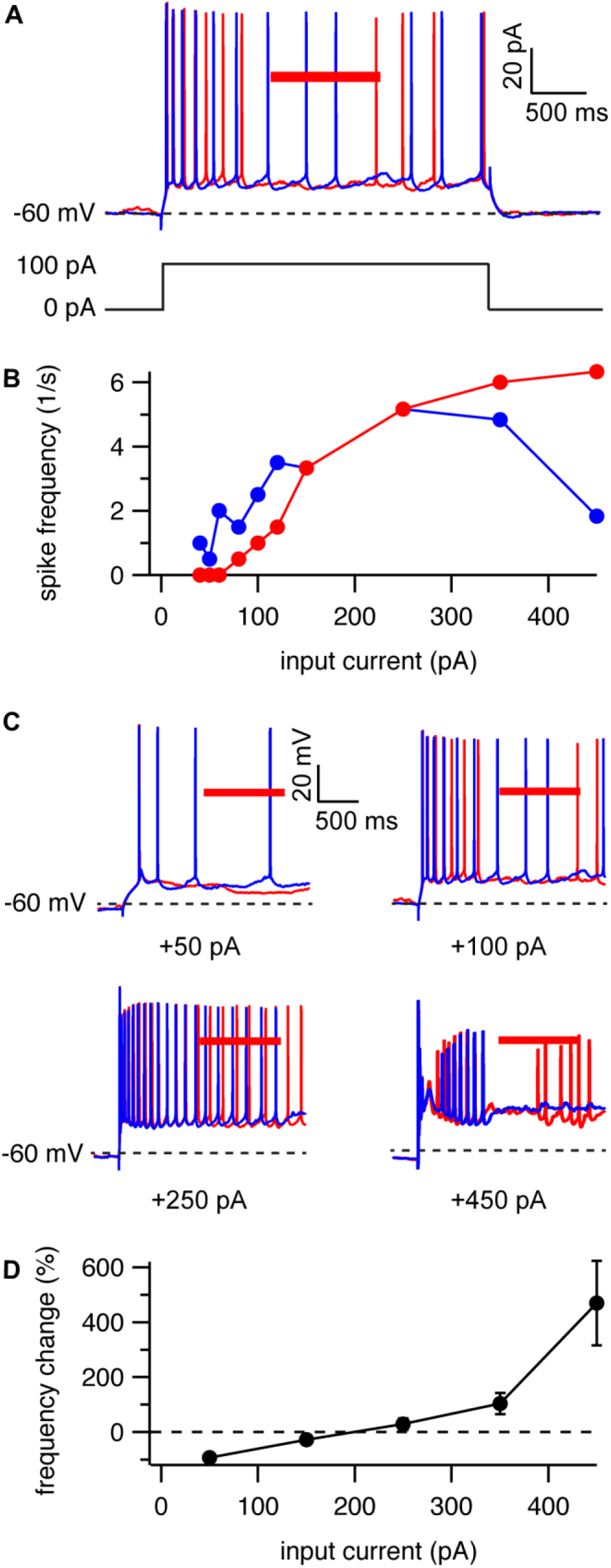
Ultrasound can also inhibit or potentiate action-potential firing when applied late in a current step. **A.** Experimental protocol and example voltage traces with (*red*) and without (*blue*) ultrasound. Ultrasound is applied 1 s after the start of a 3-s current step. The *dashed line* indicates the resting membrane voltage. **B.** Example frequency-input curve showing average firing frequency during the ultrasound application for the protocol in A (*red*) and during the same time window without ultrasound (*blue*), as a function of input current. Each point represents the average of three trials on one cell. **C.** Example voltage traces for the cell in B showing action potential firing during the ultrasound application (*red*) and during the same time window without ultrasound (*blue*) in response to current steps to either +50, +100, +250, or +450 pA. The *dashed lines* indicate the approximate resting membrane voltage of −60 mV. **D.** Mean (±SEM) change in spike frequency in response to ultrasound as a function of input current (N = 3 cells at +50 pA; N = 6 cells at all other input currents). (In 3 cells, the firing frequency at +50 pA was zero for both the control and ultrasound conditions).

Ultrasound also had a bidirectional, spike-frequency-dependent effect on excitability when it was applied 1 second after the start of a current step (Figure 3). Again, ultrasound decreased firing frequency in the low-firing-frequency, near-threshold region of the f-i curve, and increased spike rate in the high-firing-frequency, suprathreshold region of the curve (Figure 3B-D). However, the excitatory effect was more pronounced than we observed when the ultrasound pulse started 500 ms before the start of the current step. Here, ultrasound potentiated firing frequency by several hundred percent for high input currents (Figure 3D), as compared with a maximum potentiation of 49 ± 16% at 450 pA seen with the earlier ultrasound application (Figure 2C). This reflects the fact that in response to prolonged injection of high amplitude currents, accumulation of voltage-gated Na^+^ channel (Na_V_ channel) inactivation can drive pyramidal cells into a refractory state where spiking is infrequent and irregular or entirely absent (for example, the voltage trace for the control condition at +450 pA in Figure 3C shows an initial steep decline in action potential height, followed by a gradual partial recovery of action potential height, followed by a period of no action potential activity). If neurons are in this refractory state during the ultrasound application, ultrasound can “rescue” firing (as in the example voltage trace for the ultrasound condition at +450 pA in Figure 3C). This refractory state probably does not occur under normal physiological conditions, but the ability of ultrasound to rescue action-potential firing under these conditions still provides an important clue as to the molecular mechanisms underlying the effects of ultrasound on firing frequency, as discussed further below. A hint of this rescue phenomenon is also seen when the ultrasound application starts before the current step, as seen in the abrupt increase in the potentiation effect at +450 pA (Figure 2C).

### Effects of ultrasound on interspike intervals

To examine the effects of ultrasound on action-potential firing in more detail we compared the latency to the first spike, and the intervals between subsequent spikes (interspike intervals), in the control and ultrasound conditions (Figure 4). To summarize these results, and to account for the variability in intrinsic excitability between cells, we averaged *instantaneous* firing frequencies (latency and interspike intervals) across cells firing at approximately the same *average* firing frequency (either 5, 10, or 20 Hz) in the control condition at whatever input current was necessary to achieve these average firing frequencies, and at the same input current in the ultrasound condition (Figure 4A-C). We note, however, that this averaging procedure can obscure some of the details of ultrasound’s effects. At 5 Hz the effect of ultrasound applied 500 ms before the start of the current step is predominantly inhibitory (as seen in the longer average latency and interspike intervals in the ultrasound as compared with the control condition (Figure 4A), but the interval between the first and second spikes was actually shorter in the ultrasound condition than the control condition in some cells (6 out of 13 cells in this data set). This effect occurs because, even at relatively low average firing frequencies, pyramidal cells will occasionally fire “doublets” or high-frequency bursts of two action potentials, in which a second action potential is triggered by the after-depolarization of the initial action potential. When this occurs, ultrasound decreases the interval between spikes (Figure 4D). This result indicates that the mechanism by which ultrasound potentiates firing at high average firing frequencies is also active under conditions of low overall average firing frequency, during localized periods of high-frequency firing. A similar combination of inhibitory and excitatory effects can be observed at 10 Hz and even 20 Hz (Figure 4B-C). At 20 Hz, ultrasound still increased the latency to the first spike (16 ± 2 ms versus 12 ± 1 ms in the control condition; N = 12, P = 5.2 x 10^-4^, paired, two-tailed Student’s t-test), despite decreasing the interspike interval for all subsequent spikes (Figure 4C). The effect of ultrasound was also mixed when ultrasound was applied 1 s after the start of the current step (Figure 4E).

**Figure 4.**
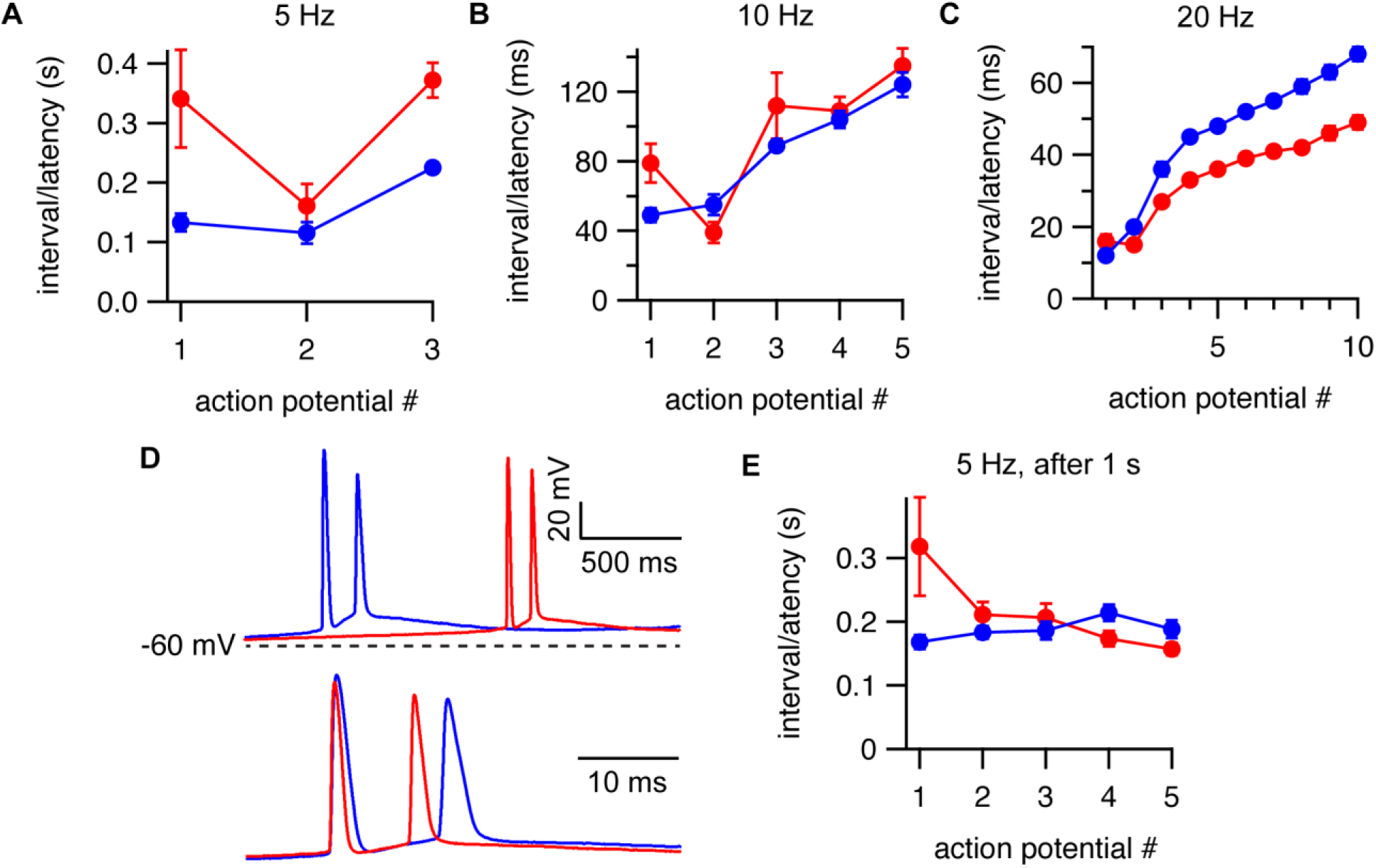
Effects of ultrasound on action potential timing. **A-C.** Mean (±SEM) latency between the start of the current step and the first action potential, and mean intervals between the first and second, and second and third, etc. action potentials, for cells firing at average frequencies of approximately 5 Hz (N = 13 cells), 10 Hz (N = 15 cells), and 20 Hz (N = 13 cells) during the first 500 ms of the current step, with (*red*) or without (*blue*) a 1-s ultrasound pulse starting 500 ms before the current step. The actual spike frequencies were 5.6 ± 0.1, 10.5 ± 0.2, and 20.4 ± 0.3 Hz, and the injected currents were 70 ± 7, 115 ± 9, and 292 ± 28 pA for the 5, 10, and 20 Hz conditions. **D.** Example voltage traces showing decreased interval between the first and second action potentials at low average firing frequency. The top panel shows the first two action potentials for the control (*blue*) and ultrasound (*red*) conditions. The *dashed line* indicates the approximate resting membrane voltage of −60 mV. The bottom panel shows the same data, aligned to the action potential threshold on a zoomed-in time scale. **E.** Same as A-C, for an average firing frequency of 5 Hz, except that ultrasound was applied 1-s after the start of a 3-s current step, and the average firing frequency and intervals/latency were determined during ultrasound stimulus or during the same time period without ultrasound (N = 6). The actual spike frequency was 5.2 ± 0.2 Hz, and the injected currents was 183 ± 21 pA.

### Effects of ultrasound on resting membrane potential

Ultrasound also has effects on resting membrane potential, which can be observed by averaging several voltage traces aligned to the onset of the ultrasound pulse, in the absence of injected current. As shown in Figure 5A, ultrasound has a slight hyperpolarizing effect on resting membrane potential. The average voltage traces also show another interesting effect of ultrasound. In addition to the relatively constant hyperpolarization, there is a transient *depolarization* of the resting membrane potential, preceding the hyperpolarization effect and acting on a faster time scale, at the onset of the ultrasound pulse; this transient depolarization is matched by a roughly symmetrical transient hyperpolarization at the offset of the pulse (Figure 5A, *arrows*). The symmetrical, on/off nature of these transients suggests that they are caused by changes in membrane capacitance, as does the fact that they occur much faster than the steady-state changes in resting membrane potential (Figure 5A-C). (Changes in membrane potential due to changes in capacitance can occur much faster than those due to ionic currents because they do not involve actual redistribution of charges across the membrane and are therefore not limited by membrane conductance). Thus, the results in Figure 5 can be described by three distinct steps: 1) ultrasound rapidly increases membrane capacitance (C) at the onset of the ultrasound pulse, which causes the membrane voltage (V) to become less negative (due to an increase in the denominator in the equation V=Q/C, where Q is the negative total charge on the membrane); 2) ionic currents then slowly change the membrane voltage to a steady-state value determined by the total ionic current (one or more ion channels having been affected by ultrasound) resulting in membrane hyperpolarization; 3) at the offset of the ultrasound pulse, capacitance rapidly relaxes back to its initial value, producing a transient decrease in membrane voltage, through essentially the same mechanism as in step 1. We investigate the physical basis of these capacitance changes and their relationship to the effects of ultrasound on excitability further below, but one point worth mentioning here is that the change in capacitance is too small for its effect on the rate of membrane charging (less than 1% change in membrane time constant) to have a significant effect on excitability in and of itself.

**Figure 5.**
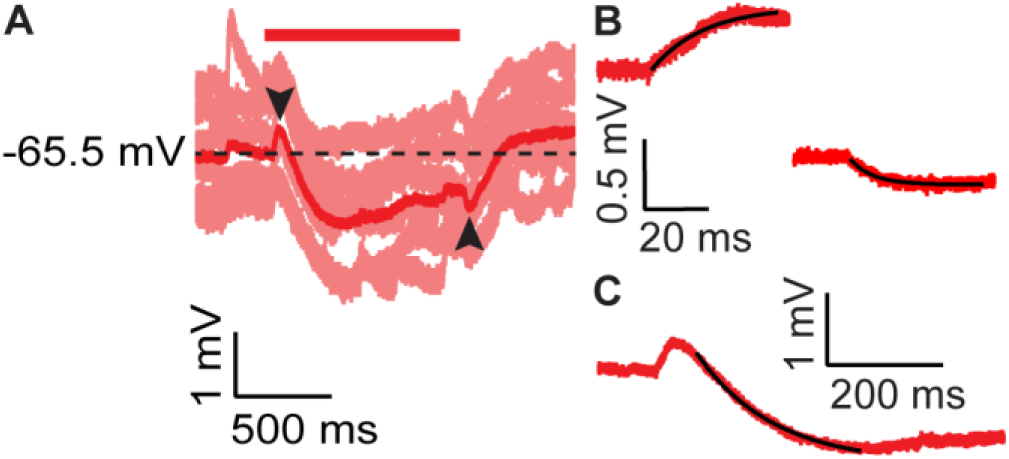
Effects of ultrasound on resting membrane potential and membrane capacitance. **A.** Six individual voltage traces (*pink*) and the average of these voltage traces (*red*) showing the effect of ultrasound (*red bar*) on resting membrane potential. The *black arrows* indicate transients due to changes in membrane capacitance. The *dashed line* indicates the mean resting membrane voltage. **B.** Zoomed-in timescale showing the fast voltage transients (*black arrows* in A) for ultrasound onset (*top left*) and offset (*bottom right*). Exponential fits (*black lines*) to the rise and fall of the voltage transients give amplitudes and time constants of 0.45 mV and 16.3 ms for the onset and 0.22 mV and 6.7 ms for the offset, for the example shown; mean values (±SEM) were 0.41 ± 0.04 mV and 10.4 ± 1.2 ms for the onset and 0.35 ± 0.04 mv and 9.7 ± 1.7 ms for the offset (N = 15). No significant differences were found between the time constants (P = 0.73) or the amplitude (P = 0.087) of the transients (paired, two-tailed Student’s t-tests) **C.** Slow membrane hyperpolarization in response to ultrasound from the average trace in panel A on a zoomed-in scale, along with an exponential fit (*black line*) to the initial hyperpolarization. The amplitude and time constant of the exponential fit were 1.55 mV and 132 ms for the example shown; mean values (±SEM) were 2.4 ± 0.3 mV and 173 ± 18 ms (N = 15).

### The K2P channel hypothesis

What ion channels might be responsible for the effects of ultrasound on excitability and resting membrane potential? A compelling hypothesis—able to explain all of our data—is that ultrasound activates a fast-activating, non-inactivating potassium channel, such as members of the K2P potassium channel family. While commonly described as “voltage-independent”, K2P channels (with the exception of TWIK-1 channels) actually have an outwardly-rectifying, voltage-dependent open probability under physiological K^+^ gradients due to the interaction of permeant ions in the selectivity filter with an activation gate (Schewe et al., 2016). Nonetheless, their rate of activation is fast (millisecond time-scale) and voltage-independent, such that they are functionally similar to truly voltage-independent channels with outwardly rectifying single-channel conductance. A primary reason for suspecting K2P channels is that the effects of ultrasound are similar regardless of whether ultrasound is presented 500 ms before or 1 s after the start of the current step (compare Figures 2 and 3), consistent with the idea that ultrasound affects a channel that does not undergo prolonged voltage-dependent inactivation during sustained depolarizations. Related to this point, the effects of ultrasound are not diminished by repetitive, high-frequency action potential firing (for example, in Figure 4C effects of ultrasound are clearly present throughout the entire 20-Hz, 10-spike train). Further, a striking feature of the effects of ultrasound on spike intervals is that ultrasound always increases the latency to the first spike (Figure 4A-C), regardless of the input current and the rate of approach to the initial action potential threshold, consistent with the idea that ultrasound affects a fast-activating, non-inactivating channel.

CA1 pyramidal neurons express a variety of K2P channel subunits. Expression of TASK-1 and TASK-3, TREK-1 and TREK-2, TRAAK, and TWIK has been shown at the mRNA level (Talley et al., 2001), while expression at the protein level has been shown has been shown for TASK-3 in CA1 pyramidal neurons specifically (Marinc et al., 2014), and for TRAAK throughout the central nervous system (Brohawn et al., 2019). In addition, functional expression of TASK-like currents has been shown in CA1 pyramidal neurons using patch-clamp recording (Taverna et al., 2005). TREK and TRAAK channels are particularly interesting in the present context since they are exceptionally sensitive to mechanical force and to increases in temperature between approximately 20 and 40°C (Maingret et al., 2000; Kang et al., 2005). Thus, TREK and TRAAK channels are responsive to the two leading candidate mechanisms by which ultrasound at 43 MHz could modulate ion channel activity.

Activation of K2P channels by ultrasound could account for all of the neurophysiological effects of ultrasound described so far: hyperpolarization of resting membrane potential; inhibition of action-potential firing in response to near-threshold current injections; and— although this last point may seem counter-intuitive—potentiation of action potential firing at high firing frequencies. Hyperpolarization of resting membrane potential by increased outward K^+^ current is straightforward, as is the idea that K^+^ current can inhibit firing, but K^+^ current can also potentiate firing by its effects on action-potential repolarization and afterhyperpolarization (AHP). By accelerating the rate of membrane repolarization following the peak of an action potential (thereby reducing action potential width) K^+^ current can reduce inactivation of voltage-dependent Na^+^ channels during the action potential, and by increasing the depth of the AHP, it can accelerate the voltage-dependent recovery of Na_V_ channels from inactivation. Both of these effects would tend to increase the population of Na_V_ channels available for activation in response to depolarizing current and would increase the maximum action potential firing frequency, as we in fact see in response to ultrasound (Figures 2 and 3). This mechanism is well-known and widespread in neurophysiology, with several K^+^ channels, including both K2P channels and voltage-dependent K^+^ channels (K_V_ channels), having been shown to facilitate high-frequency firing (Lien and Jonas, 2003; Brickley et al., 2007; Gu et al., 2007; Gonzalez et al., 2009; Liu and Bean, 2014; Kanda et al., 2019). The idea that the potentiation of firing by ultrasound is due to effects on action-potential repolarization and AHP is supported by the results in Figure 4C. For neurons firing at an average firing frequency of 20 Hz in the absence of ultrasound, ultrasound increases the latency to the first spike, while it decreases the intervals between all subsequent spikes. The lack of a potentiating effect on the first spike, despite the otherwise strongly potentiating effects of ultrasound, indicates that the potentiating effect acts through a process (such as action-potential repolarization and AHP) that occurs *after* the initiation of the first action potential. The idea that ultrasound can potentiate firing by activating K^+^ current makes specific predictions about the effects of ultrasound on action potential waveform: ultrasound should accelerate repolarization, decrease action potential width, and increase the depth of the AHP.

### Effects of ultrasound on action-potential waveform

To test the idea that potentiation of firing by ultrasound is due to activation of K^+^ channels, we examined the effects of ultrasound on action-potential waveform in our recordings. The effects of ultrasound on action-potential waveform are consistent with the idea that ultrasound facilitates high-frequency firing by accelerating action-potential repolarization. Figure 6A-F shows the effect of ultrasound applied 500 ms before the start of the current step on action-potential width for cells firing at average frequencies of 5, 10, and 20 Hz. (Ultrasound also had effects on action-potential height, although these were less pronounced than the effects on width; effects on height are detailed in Supplemental Figure 3.) Ultrasound decreased action-potential width for every action potential at all firing frequencies. As shown in Figure 6G-H, ultrasound also decreased action-potential width when applied 1 s after the start of a current step, again indicating that the channels responsible for these effects continue to influence firing frequency and remain responsive to ultrasound throughout sustained depolarizing current steps.

**Figure 6.**
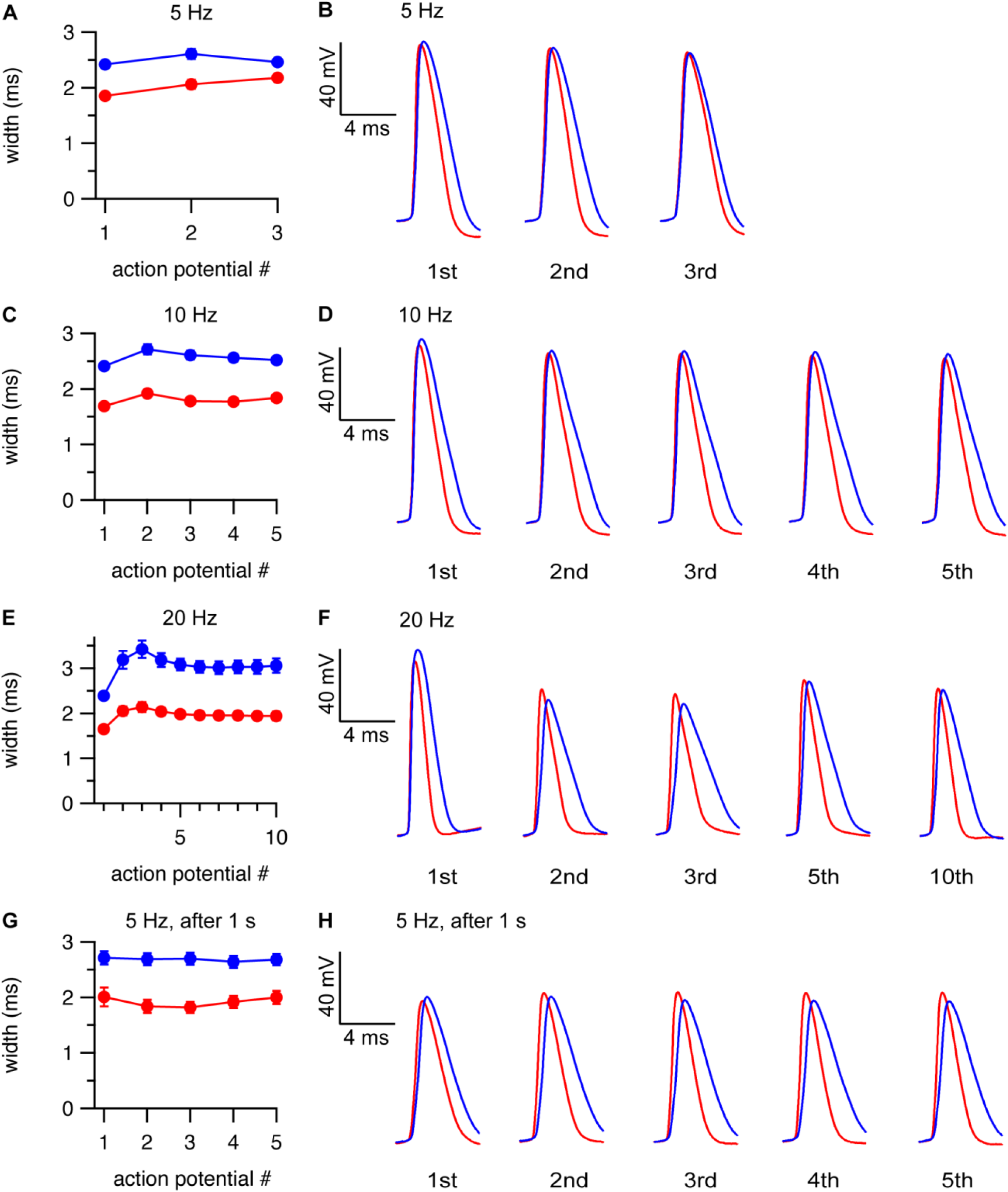
Ultrasound decreases action-potential width. **A-B.** Mean (±SEM, N = 13 cells) action-potential widths (A) and example action-potential waveforms aligned to the action-potential threshold (B) as function of action-potential number in the presence (*red*) and absence (*blue*) of a 1-s ultrasound pulse starting 500 ms before the current step, for cells firing at an average firing frequency of approximately 5 Hz (as measured during the first 500 ms of the current step) in the control condition. **C-D.** As in A-B, but for cells firing at an average firing frequency of approximately 10 Hz in the control condition (N = 15). **E-F.** As in A-B, but for cells firing at an average firing frequency of approximately 20 Hz in the control condition (N = 13). **G-H.** As in A-B, but with ultrasound applied 1 s after the start of a 3-s current step, for cells firing at an average firing frequency of approximately 5 Hz in the control condition, with firing frequency determined in a 1-s window starting 1 s after the current step (corresponding to the time period of the ultrasound stimulus), and action potential number relative to the start of the ultrasound stimulus (N = 6).

The effects of ultrasound on action-potential width tended to counteract the broadening of action-potential width that occurs during high-frequency firing. Figure 7 plots action potential width as a function of action potential number and input current for the control and ultrasound conditions. In the control condition, there are dramatic differences in width between the first action potential and subsequent action potentials at high input currents, while in the ultrasound condition these differences are much less pronounced. To quantify this effect, we measured the difference in width between the first and third, first and fifth, and first and last action potentials during the ultrasound stimulus, in response to a +450 pA current step for the control and ultrasound conditions. These differences (mean ± SEM, control vs. ultrasound, N = 9) were 2.6 ± 0.9 vs. 0.7 ± 0.1 ms, 1.5 ± 0.1 vs. 0.5 ± 0.1 ms, and 1.2 ± 0.2 vs 0.5 ± 0.1 ms for the third, fifth, and last action potentials (P = 0.064, 1.4 x 10^-5^, and 0.0013, paired, two-tailed Student’s t-test). One plausible interpretation of this result is that K2P channels activated by ultrasound cause the membrane voltage during the action potential to repolarize before slower-activating K_V_ channels, which would otherwise contribute to action-potential repolarization, are activated. Since time-dependent activation and inactivation of K_V_ channels causes action-potential broadening during repetitive firing (Giese et al., 1998; Shao et al., 1999; Yue and Yaari, 2004; Kim et al., 2005; Gu et al., 2007), an increase in the contribution of K2P channels lacking time-dependent inactivation with a concomitant decrease in the contribution of K_V_ channels to action-potential repolarization would reduce time- and frequency-dependent action potential broadening.

**Figure 7.**
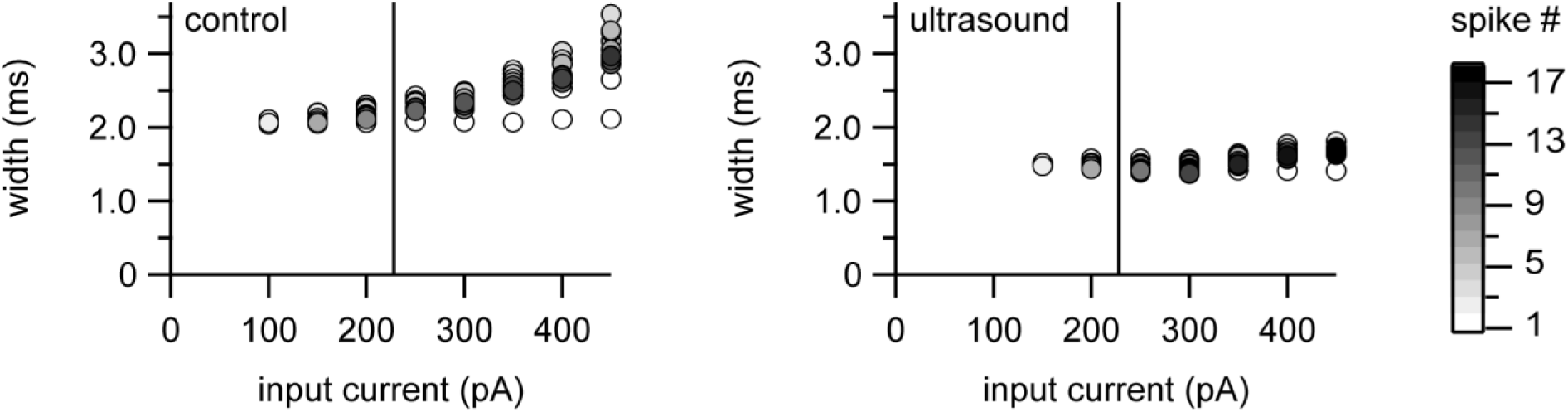
Ultrasound reduces action-potential broadening. Example data showing action-potential width as a function of action-potential number (indicated in *grayscale*, scale bar at far left) and input current level, for the first 500 ms of the current step, for currents from 0 to +450 pA in 50 pA steps, with (*right*) or without (*left*) a 1-s ultrasound pulse starting 500 ms before the current step. The vertical lines indicate the approximate location of the transition between inhibitory and potentiating effects of ultrasound.

Although the effects of ultrasound on action-potential widths are predominantly due to acceleration of the repolarization phase, we also noted effects on the rising phase of the action potential. These effects are readily apparent in the first derivative of membrane voltage during the action potential (Figure 8A) or in action potential phase plots (plots of the first derivative of voltage versus voltage, Figure 8B). In fact, the maximum rates of voltage rise and fall during the action potential were both consistently increased by ultrasound throughout a spike train (Figure 8C-D). The effect on the falling phase is to be expected based on the observed decrease in spike width and the hypothesis that ultrasound activates K2P channels. The effect on the rising phase is also consistent with this hypothesis, as activation of K2P leading to reduced Na_V_ channel inactivation would increase the number of Na_V_ channels available to activate during the rising phase of the action potential. Alternatively, increased K2P conductance could increase the rate of action potential rise by decreasing the membrane time constant (Brickley et al., 2007). Consistent with these results, decreases in the rates of action-potential rise and fall were seen with knock-outs of K2P channels in cerebellar granule neurons (Brickley et al., 2007) and hypothalamic hypocretin/orestin neurons (Gonzalez et al., 2009), while in a heterologous action-potential firing model higher levels of K2P expression increased the rate of action-potential rise (MacKenzie et al., 2015).

**Figure 8.**
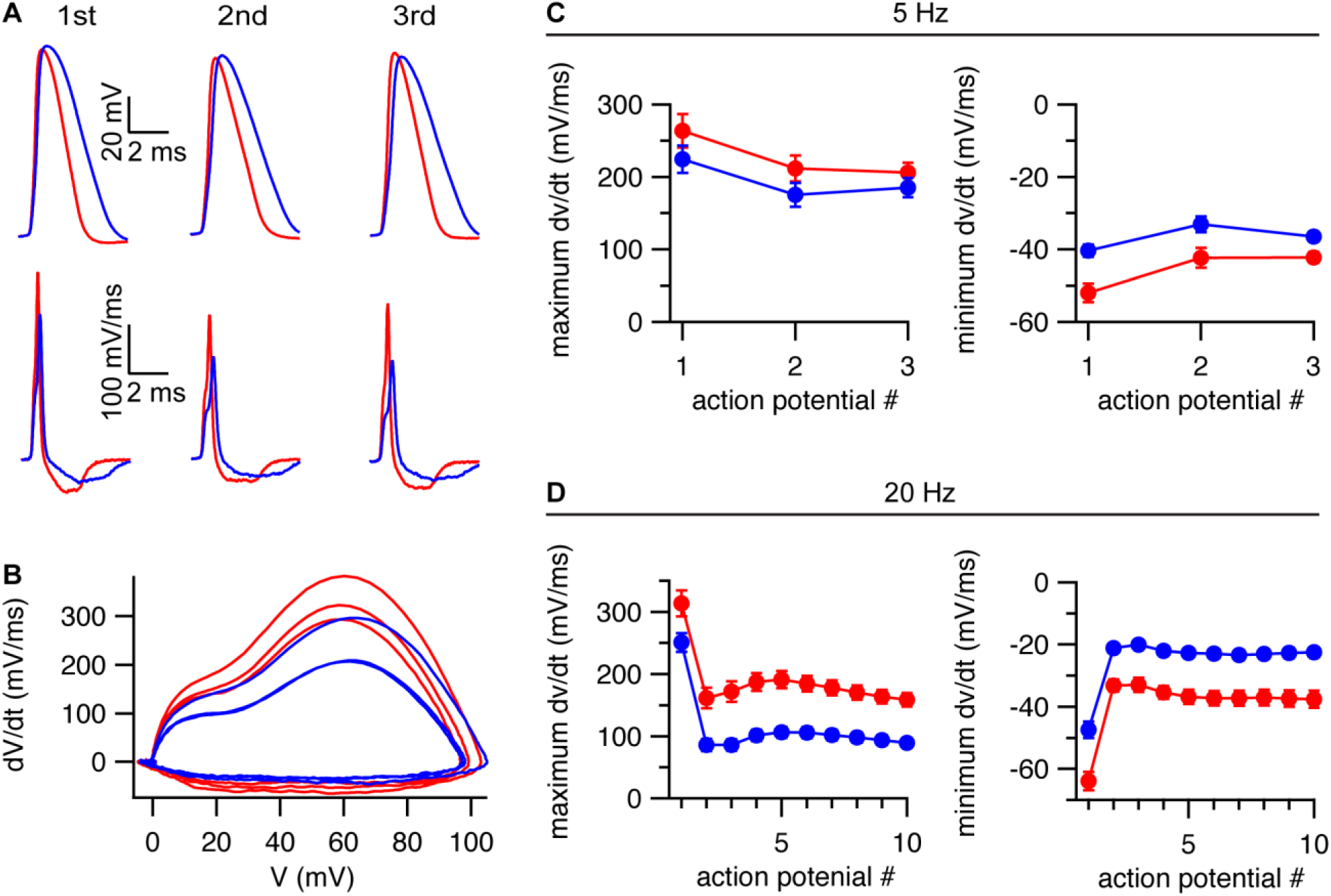
Effects of ultrasound on depolarization and repolarization rates. **A.** Example traces showing the membrane voltage (*top*) and its first derivative (*bottom*) for the first three action potentials in response to a +100 pA current step in the presence (*red*) and absence (*blue*) of a 1- s ultrasound pulse starting 500 ms before the current step, aligned to the action potential threshold. **B.** Phase plots for the action potentials shown in panel A. **C-D.** Maximum rates of depolarization (*left)* and repolarization (*right*) during the action potential (mean ± SEM, N = 13 cells), as a function of action potential number, in the presence (*red*) and absence (*blue*) of a 1-s ultrasound pulse starting 500 ms before the current step, for cells firing at an average firing frequency of 5 Hz (**C**) or 20 Hz (**D**) in the control condition.

In addition to effects of ultrasound on the rising and falling phases of the action potential, we also found effects on the AHP. To quantify these effects, we measured the voltage minimum between action potentials during repetitive firing. Because measurements of this parameter are very sensitive to changes in resting membrane voltage and series resistance that can occur over long recording times, we compared voltage minimums before, during, and after ultrasound application within the same voltage trace (Figure 9A-B) and made a similar comparison for control recordings. We performed these comparisons for ultrasound applied 1 s after the start of a current, for cells firing at an average frequency of 5 Hz in the control condition. This firing frequency is near the transition region between the inhibitory and potentiating effects of ultrasound on spike frequency, such that the spike frequency is similar for the control and ultrasound conditions, allowing us to compare a similar number of interspike voltage minima for the control and ultrasound conditions (Figure 9C). This analysis demonstrates that the depth of the AHP is greater during the ultrasound application than before or after it, or during the same time windows for the control condition. Together with the effects of ultrasound on spike waveform (Figure 6), this result supports the idea that ultrasound activates a sustained outward current, which limits Na_V_ channel inactivation and thereby potentiates high-frequency firing. Removal of Na_V_ channel inactivation by membrane hyperpolarization also explains how ultrasound can rescue spiking in neurons that have entered a refractory state due to accumulation of Na_V_ channel inactivation (Figure 3C, *bottom right*).

**Figure 9.**
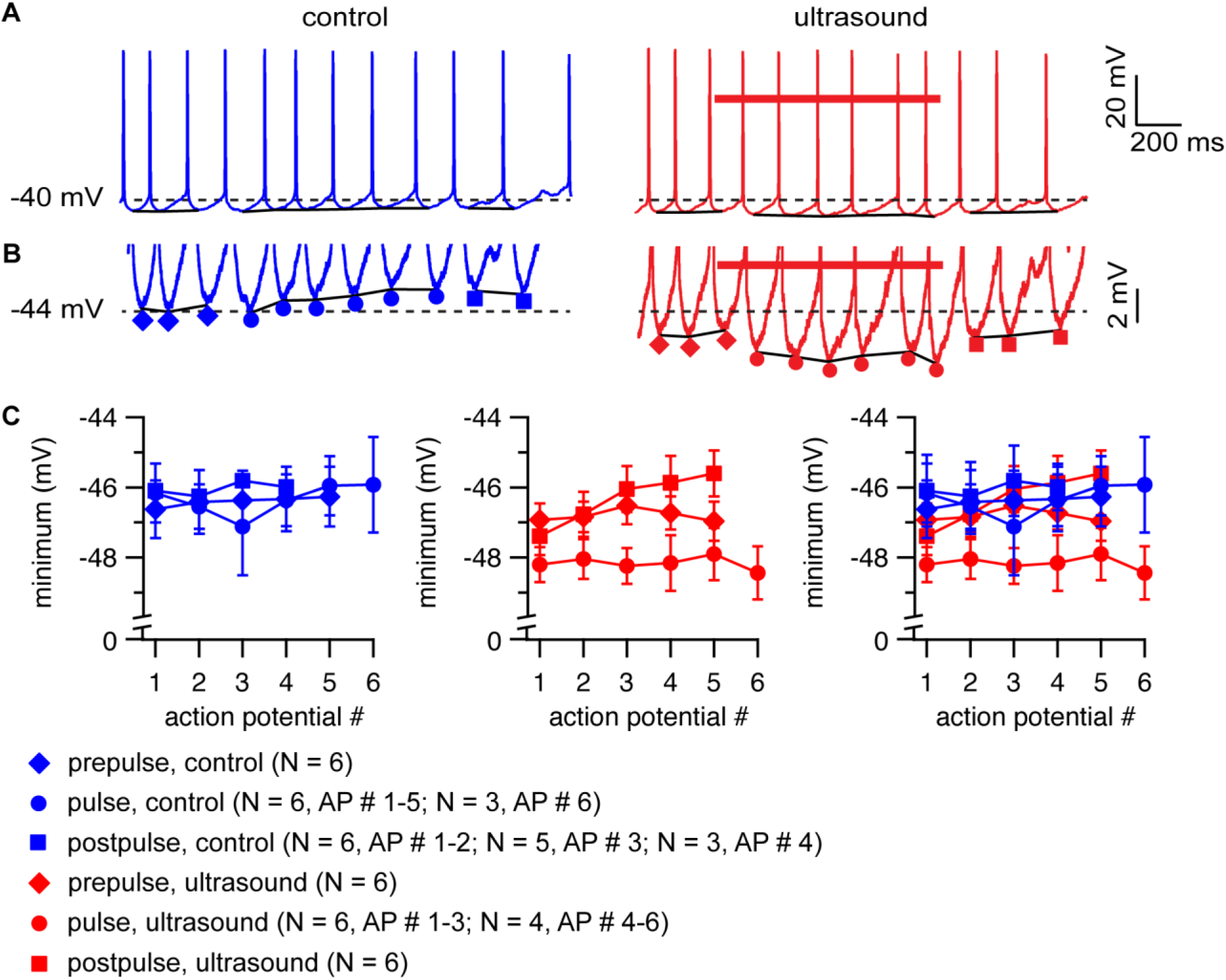
Ultrasound increases the depth of the afterhyperpolarization. **A.** Example voltage traces comparing voltage minima between action potentials in response to a 3-s, 250-pA current step with (*red voltage trace, right*) and without (*blue voltage trace, left*) a 1-s ultrasound pulse (*red bar*) starting 1 s after the start of the current step. The *solid black lines* connect the voltage minima between action potentials before, during, or after the ultrasound pulse, or during the corresponding time periods for the control condition. The dashed lines indicate a reference voltage level of −40 mV. The resting membrane voltage for this cell was −63 mV. **B.** Same as panel A, on a zoomed-in voltage scale. The *red diamonds* indicate voltage minima before the ultrasound pulse, *red circles* indicate voltage minima during the ultrasound pulse, and *red squares* indicate voltage minima following the ultrasound pulse; *blue symbols* indicate voltage minima for the corresponding time periods for the control condition. The dashed lines indicate a reference voltage level of −44 mV. **C.** Mean (±SEM, N = 3-6 cells, see figure panel for details) values of the voltage minimum, as a function of action potential (AP) number for the first four to six action potentials before, during, and after the ultrasound pulse, along with the equivalent mean values for the control condition, following the symbolism indicated in panel B. The means were determined for cells firing at the same average frequency (5 Hz) during a 1-s window starting 1 s after the start of the current step (corresponding to the period of the ultrasound stimulus) in the control condition. For clarity, the results are shown separately for the control group only (*left*), the ultrasound group only (*middle*), and for both groups simultaneously (*right*). Significant differences between groups were only found in the presence of ultrasound (P = 1.0 x 10^-4^, 1.4 x 10^-5^, and 0.16 for before versus during, during versus after, and before versus after the ultrasound pulse; P = 0.92, 0.39, and 0.35 for comparisons of the same time periods in the control condition (unpaired, two-tailed Student’s t-tests, unequal variance).

### Physical mechanism of neuromodulation by high-frequency ultrasound

The idea that ultrasound acts on K2P channels is also consistent with the physical effects of ultrasound on biological tissue. At 43 MHz, two plausible mechanisms through which ultrasound might modulate the activity of ion channels are heating and mechanical stress due to acoustic radiation force. Absorption of acoustic energy by biological tissue as heat can increase its temperature, with effects on ion channel gating and all other biological reactions. Absorption also results in attenuation of ultrasound intensity as the wave propagates, creating spatial gradients in intensity that give rise to radiation force, which in turn produces tissue displacement and strain. At the microscopic scale, this displacement and strain may involve increased tension in the cell membrane, cytoskeleton, and extracellular matrix, all of which may affect excitability through mechanical effects on ion channel proteins. Among the K2P channels that may be expressed by CA1 pyramidal cells, TREK and TRAAK channels are especially sensitive to thermal and mechanical stimuli.

To gain further insight into these physical mechanisms, we performed finite-element simulations of the effects of ultrasound on brain slices in the context of our experimental set-up. The simulated spatial profiles of ultrasound-induced heating and macroscopic tissue displacement in response to radiation force are shown in Figure 10A-B. Notably, the spatial profiles of heating and displacement effects are significantly wider than the 90-micron diameter of the focal volume of the ultrasound beam, with significant heating and displacement occurring several hundreds of microns away from the beam axis (Figure 10C). This is an important result because the thermo- and mechanosensitive K2P channels TREK-1 and TRAAK are expressed at high density at the nodes of Ranvier of vertebrate neurons (Brohawn et al., 2019; Kanda et al., 2019). The first node of Ranvier is located approximately 100 microns from the axon initial segment (Kole, 2011). In our experiments, the soma of the patched neuron is located approximately on the axis of the ultrasound beam (see *Materials and Methods*) so the first node of Ranvier is probably within the region of the tissue exposed to thermal and mechanical effects of ultrasound. Axonal K^+^ channels play important roles in regulating excitability (Shah et al., 2008; Kole, 2011; Kanda et al., 2019). It is therefore plausible that a subpopulation of TREK-1 or TRAAK channels at the nodes of Ranvier could contribute to the neurophysiological effects of ultrasound. The magnitude of the temperature change is also consistent with a thermal mechanism for the effects of ultrasound. The maximum temperature change in the simulation is 1.3 C; temperature changes of this size have previously been shown to affect neural excitability (Owen et al., 2019). The maximum value of the simulated displacement (1.7 microns) is similar to the displacement measured in the retina during ultrasound neuromodulation with stimulus parameters similar to those used here (Menz et al., 2019).

**Figure 10.**
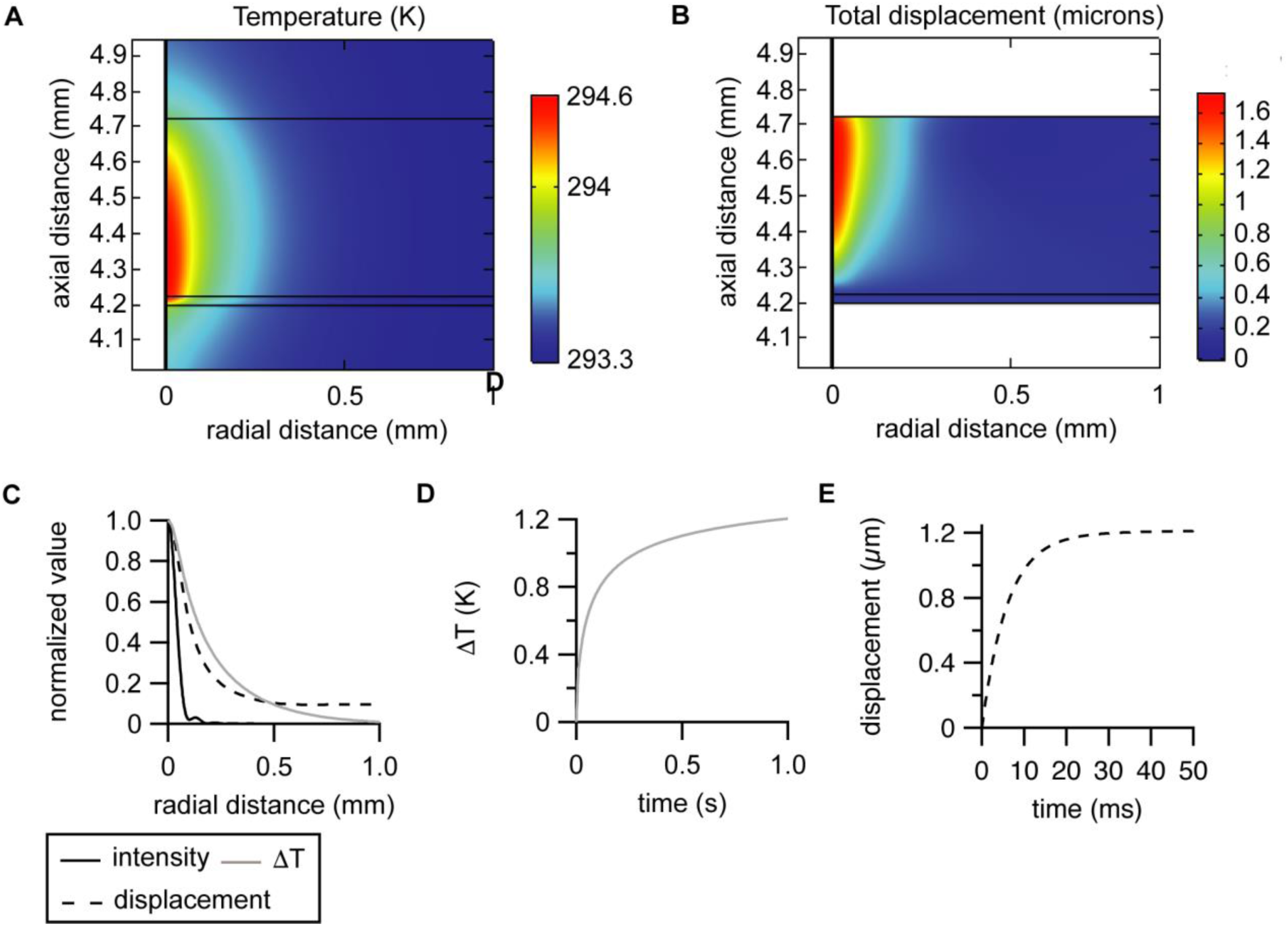
Simulated tissue heating and displacement in response to ultrasound. **A.** Spatial profile of temperature after one second of ultrasound exposure at 43 MHz and 50 W/cm^2^, as a function of axial distance from the transducer surface and radial distance from the ultrasound beam axis, in a 500-micron thick brain slice and 25-micron thick polystyrene film (*middle two layers*) and the surrounding fluid (external solution, *top layer*, and distilled water, *bottom layer*). **B.** Spatial profile of the static total displacement in response to acoustic radiation force in the brain slice and polystyrene film. **C.** Normalized values of the acoustic intensity (*solid black line*), temperature rise after 1 second of ultrasound exposure (*solid gray line*), and total displacement (*dashed line*), at a depth of 250 microns in the brain slice, as a function of radial distance. **D.** Time course of the temperature rise in response at ultrasound at a depth of 250 microns in the brain slice on the axis of the ultrasound beam. The time course of the temperature change can be described by two exponential components with amplitudes and time constants of −0.56 C and 30 ms, and −0.57 C and 295 ms, for a weighted time constant of 164 ms. **E.** Time course of the displacement in response at ultrasound at a depth of 250 microns in the brain slice on the axis of the ultrasound beam. The time course of the displacement change can be described by two exponential components with amplitudes and time constants of −1.2 microns and 6 ms, and −0.13 microns and 344 ms, for a weighted time constant of 39 ms. Note that the steady-state displacement is slightly smaller than in the static displacement simulation due to the inclusion of the fluid loading in the dynamic displacement simulation.

Since the activity of TREK and TRAAK channels is highly temperature-sensitive, we considered whether our results might represent an artefact due to the experiments being performed at room temperature (21-23°C). Room temperature is near the threshold for temperature activation of these channels, such that they are mainly inactive in the absence of other stimuli such as membrane tension, lipid agonists, or acidic pH (Maingret et al., 2000; Kang et al., 2005). Thus at room temperature in the absence of additional gating stimuli, the relative increase in thermosensitive K2P current would be greater in our experiments than at physiological temperatures, which might lead us to overestimate the importance of K2P channels in the response to ultrasound. On the other hand, the midpoints of the temperature-activation curves for TREK and TRAAK channels are near 37°C (in other words, near body temperature in mammals) so that the absolute increase in K2P current in response to increased temperature is near maximal at physiological temperatures, so we might instead be underestimating the thermal effects of ultrasound on K2P channels that would occur in physiological contexts. To cut through this speculation and address these issues, we repeated our experiments measuring the neurophysiological effects of ultrasound in cells firing at a spike frequency of approximately 5 Hz at near-physiological temperature (30°C) (Supplemental Figure 4). Similar to what we observed at room temperature, ultrasound inhibited action potential firing at this relatively low spike frequency and decreased action potential width, indicating that these effects are not especially sensitive to the ambient temperature. However, the hyperpolarizing effect of ultrasound on the resting membrane voltage that we observed at room temperature was no longer apparent at 30°C, possibly due to increased noise in the baseline voltage or more hyperpolarized (in other words, closer to the K^+^ reversal potential) resting voltage at higher temperature.

It is instructive to consider the amplitude and time course of the membrane capacitance change in response to ultrasound (Figure 5) in the context of possible thermal and mechanical mechanisms. As described above (under *Effects of ultrasound on resting membrane potential*), the capacitance change in response to ultrasound is fast relative to the membrane time constant, so we can assume that the total charge on the membrane is constant during the initial transient depolarization in response to ultrasound (Figure 5A, *left arrow*). In other words, the numerator in the equation V = Q/C is constant, so for small changes in voltage the relative change in voltage is approximately inversely proportional to the relative change in capacitance. Empirically, membrane capacitance increases by approximately 1% per degree C (Taylor, 1965). This is consistent with the size of the simulated temperature rise (peak simulated temperature rise of 1.3°C compared with the measured amplitude of the initial decrease in voltage of 0.7 ± 0.1%). However, the time course of the temperature rise (Figure 10D) is much slower than the time course of the voltage transient (which again, assuming constant Q, is identical to the time course of the capacitance change). The time course of the change in resting membrane potential (173 ± 18 ms, Figure 5), however, parallels that of the temperature rise. (A capacitance change on the time course of the simulated temperature rise would not have a significant effect on the membrane voltage, as it would be counteracted by ionic currents.) Thus, the simulated ultrasound heating results strongly suggest that ultrasound affects action-potential firing in our experiments at least in part through a thermal effect on ion channels, but do not explain the presence of the capacitive transients.

We considered whether the time course of the capacitive transients could instead be explained by the dynamics of the tissue mechanical response to acoustic radiation force. We sought to determine whether, having already modeled the static displacement of the tissue, we could, without retroactively changing any of the tissue material properties, obtain a time course for tissue displacement similar to that of the capacitive transients using a simple viscoelastic model with reasonable tissue viscous properties (see *Materials and Methods*). We found that this could be achieved using a shear viscosity (μ) of 1 Pa·s (Figure 10E). Since the tissue is essentially incompressible in our model (Poisson’s ratio (ν) = 0.4998), this is equivalent to a relaxation time of 2μ·(1+ ν)/E = 6 ms (where E is Young’s modulus). Biological tissue is a highly heterogeneous material that displays a variety of active and passive mechanical responses to force, spanning time scales from milliseconds to hours (Ricca et al., 2013), and as a result its viscous properties are highly sensitive to the time scale of the measurement and even complex viscoelastic models encompassing multiple relaxation times may not fully describe the viscoelastic behavior of tissue. Nonetheless, the shear viscosity/relaxation time in our model is reasonable for a soft, gel-like material, and is comparable to fast relaxation times observed experimentally in brain tissue (Arbogast and Margulies, 1999; Abolfathi et al., 2009; Rashid et al., 2012, 2013). Moreover, the simulated time course of displacement is consistent with experimental measurements of the tissue displacement in response to ultrasound at 43 MHz and 40 W/cm^2^ in the salamander retina, which was found to be complete in less than 10 ms (Menz et al., 2019).

We can therefore make the reasonable assumption that the capacitive transients are due to a mechanical effect on membrane properties, and we can estimate the size of the potential ion-channel gating effects that would occur as a result of this mechanical effect. The capacitance of a lipid bilayer membrane is given by C = ε·ε_0_·A/L, where ε is the dielectric constant of the hydrophobic core of the lipid bilayer, ε_0_ is the permittivity (polarizability) of free space, A is membrane area, and L is the thickness of the hydrophobic core of the membrane. For small strains like those under consideration here, lipid bilayer membranes can be considered incompressible, such that a 1% increase in capacitance corresponds to a 0.5% increase in area and a 0.5% decrease in thickness (White and Thompson, 1973; Alvarez and Latorre, 1978). An increase in membrane area can be converted to an increase in membrane tension (γ) according to γ = ΔA·K_A_, where ΔA is the relative change in area and K_A_ is the area elastic constant of the membrane. Area elastic constants measured for lipid membranes are on the order of 100’s of mN/m (Evans et al., 1976; Kwok and Evans, 1981; Needham and Nunn, 1990). If the capacitance transients are due to membrane strain, the resulting membrane tension is on the order of a few 0.1 mN/n to a few mN/m. These values are similar to the tension thresholds for activation of mechanosensitive K2P channels (estimated as 0.5-4 mN/m for activation of TREK-1 and TRAAK (Brohawn et al., 2014)), which are low relative to other known mammalian mechanosensitive channels. Notably, a recent *in vivo* ultrasound neuromodulation study of the murine sciatic nerve at 4 MHz, found that tissue displacement *in vivo* was highly correlated with the neuromodulation effects (Lee et al., 2020). Nonetheless, additional data or theoretical advances would be required to firmly associate these capacitive transients with changes in membrane tension. If such an association could be made, it would provide strong evidence that ultrasound modulates action potential firing through mechanical effects of radiation force in our experiments. At present, our results do not rule out this idea, but the case for mechanical effects remains speculative, while the role of thermal effects seems highly plausible. Nonetheless, our simulation results support the conclusion that both inhibitory and excitatory effects of high-frequency ultrasound on action-potential firing are due to activation of thermo- and mechanosensitive K2P channels.

## DISCUSSION

To summarize, the neurophysiological effects of ultrasound that we have described here can all be explained by activation of a sustained outward current. We argue that the molecular basis of this outward current is most likely one or more of the K2P channels expressed by CA1 pyramidal neurons. Although a variety of voltage-dependent K^+^ currents shape the action-potential waveform and regulate excitability in these neurons, several arguments suggest that K2P channels are the molecular basis of the ultrasound-activated outward conductance. First, the K2P channels TREK and TRAAK, being strongly mechanosensitive and thermosensitive, have biophysical properties that make them especially sensitive to physical effects of ultrasound. Second, the fact that ultrasound has similar effects on firing frequency whether it is applied 500 ms before or 1 s after the start of a current step suggests that ultrasound affects firing through a channel that does not undergo prolonged voltage-dependent inactivation during sustained depolarizations. Finally, the neurophysiological effects of ultrasound are, strikingly, essentially the opposite of those caused by knock-out of K2P channels in other neurons, as detailed in the following paragraph.

Knock-out of TASK-3 channels in cerebellar granule neurons increased excitability at low input currents, but decreased excitability at high input currents and led to failure of sustained high-frequency firing (Brickley et al., 2007). In addition, knock-out of TASK-3 decreased the maximum firing frequency and decreased action-potential height while increasing action potential width through decease in the rates of both action-potential rise and fall. Similarly, double knock-out of TASK-1 and TASK-3 in hypothalamic hypocretin/orexin neurons inhibited high-frequency action potential firing, reduced the rates of action-potential rise and fall, and decreased the depth of the AHP (Gonzalez et al., 2009). The connection between our results and these knock-out studies is supported by experiments in a heterologous model system consisting of HEK cells transfected with TREK-1 and TASK-3 (with endogenous K_V_ channels blocked) and Na_V_ channels simulated by dynamic clamp (MacKenzie et al., 2015). In this model system, high levels of K2P expression were necessary for repetitive action potential firing, and increased K2P conductance increased the rates of action-potential rise and fall and increased the threshold current for action-potential firing. (In the context of this model system, a 74% potentiation of the K2P conductance by halothane produced effects on the rates of rise fall of the same order as we see here, potentially providing an estimate of the potentiation of K2P conductance by ultrasound in our experiments. However, caution should be used in extrapolating from this heterologous model system to neurons expressing a considerably more complex array of ion channels.) Finally, TREK-1 and TRAAK channels are also necessary for high frequency firing at the nodes of Ranvier of afferent neurons (Kanda et al., 2019).

Ultrasound neuromodulation has been studied in vertebrate axon preparations, where it has generally been found that ultrasound inhibits action-potential conduction, with the effect specifically attributed to heating in some cases (Young and Henneman, 1961; Mihran et al., 1990; Tsui et al., 2005; Colucci et al., 2009). These results are consistent with the idea that activation of K2P channels by ultrasound can inhibit action potential firing, but it would be worthwhile to revisit these experiments to see whether the bidirectional, spike-frequency-dependent effect that we observe is also present in such preparations. Our results also suggest an approach to ultrasound neuromodulation in which action potential propagation is the locus of neuromodulation, targeting white-matter tracts instead of soma-dense gray matter. The idea that ultrasound-activated K^+^ currents can both inhibit and potentiate firing might also help explain why ultrasound can both inhibit and potentiate neural activity *in vivo* (Min et al., 2011).

Although activation of K2P channels is sufficient to explain our results, we do not rule out the possibility that ultrasound affects other channels in addition to K2P’s; indeed, we think it is likely that ultrasound does affect other channels to some extent. All ion-channel gating reactions are sensitive to temperature, with typical Q10 values of ∼3, such that their rates would be expected to increase by about 10% based on the temperature changes in our simulations (Hille, 2001). Mechanical effects of radiation force could also affect channels besides K2P channels. The mechanically gated channel Piezo2 is expressed in a subset of CA1 pyramidal neurons (Wang and Hamill, 2020). Piezo1 (Qiu et al., 2019) and TRP channels (Oh et al., 2020; Yoo et al., 2020) have been experimentally linked to ultrasound neuromodulation effects. In addition, most ion channels and membrane proteins, while not functioning physiologically as mechanoreceptors, are sensitive to mechanical stimuli to some extent, either through the energetics of their interactions with hydrophobic core of the lipid bilayer or through mechanical interactions with the cytoskeleton or extracellular matrix. In fact, gating of voltage-dependent Na^+^ (Morris and Juranka, 2007), K^+^ (Tabarean and Morris, 2002; Laitko and Morris, 2004; Beyder et al., 2010), and Ca^2+^ (Calabrese et al., 2002) channels, and of NMDA receptor channels (Kloda et al., 2007), can be modulated by membrane stretch in membrane patches. However, we previously were unable to detect any mechanical modulation of heterologously expressed NaV1.2 channels by ultrasound at 43 MHz and 90 W/cm^2^ under conditions where ultrasound activated the mammalian mechanoreceptor channel Piezo1 (Prieto et al., 2018). Neural Na_V_ channels and KCNQ channels interact with the periodic actin cytoskeleton of axons through spectrin and ankyrin-G at the axon initial segment and nodes of Ranvier and (Zhou et al., 1998; Pan et al., 2006; Leterrier, 2018), suggesting that they may be sensitive to modulation by cytoskeletal tension due to acoustic radiation force. The concentration of TREK-1 and TRAAK channels at the nodes of Ranvier suggest that some similar interaction with the cytoskeleton may be involved in the localization of these channels, although this has not been demonstrated, and the intracellular domains that would facilitate such interactions are relatively small in K2P channels. Ultrasound has been shown to cause changes in cytoskeletal structure (Mizrahi et al., 2012), which might also affect the activity of cytoskeleton-associated channels. Thus investigation of the role of other ion channels in ultrasound neuromodulation should continue. Notably, several of the channels discussed above have roles in synaptic transmission, so effects of ultrasound on these channels would not be revealed by our experiments on somatic excitability in response to injected current.

It is well established that high-intensity light at infrared and shorter wavelengths can modulate neural activity through tissue heating. As a general principle, thermal neuromodulation effects in response to optical stimulation would be expected to be very similar to thermal neuromodulation effects caused by ultrasound, and studies of optical neuromodulation could therefore provide useful guidance in interpreting our results. However, optically based thermal neuromodulation experiments are highly heterogeneous in terms of the neuromodulation effect, the temperature rise required to produce the effect, and the mechanistic interpretation of the results (Wells et al., 2007; Richter et al., 2011; Shapiro et al., 2012; Duke et al., 2013; Walsh et al., 2016; Lothet et al., 2017; Paris et al., 2017; Owen et al., 2019; Zhu et al., 2019), so their usefulness is limited in this respect. Both inhibition and potentiation of firing have been reported, and the increase in temperature has varied considerably, ranging from less than 1 C to 10’s of degrees C. The temperature rise in our simulations is on the low end of this range. However, it has been proposed that spatial or temporal gradients in temperature, rather than the absolute temperature change, may determine the response to thermal stimuli (Wells et al., 2007; Paris et al., 2017). In addition, heating can cause phase changes in lipid bilayers, which have been shown to modulate ion channel activity (Seeger et al., 2010); in this case, the response to heating would also depend critically on the initial temperature. Interestingly, a recent study demonstrated inhibition of firing by small (2°C or less) increases in temperature in several different types of neurons, but *not* in CA1 pyramidal neurons (Owen et al., 2019). These effects were attributed to inward rectifier (K_ir_) potassium channels, which are not expressed in CA1 pyramidal neurons. However, our results suggest that the lack of an effect in pyramidal neurons could also be explained if the experiments were performed at a point on the f-i curve where competing inhibitory and excitatory effects of heat-activated K^+^ current result in no net effect on firing frequency. A recent *in vivo* ultrasound neuromodulation study in a rat model using ultrasound at 3.2 MHz with exceptionally long ultrasound exposure times (10’s of seconds; see below for a more general discussion of *in vivo* ultrasound neuromodulation studies) also found that inhibition could be produced by ultrasound-induced temperature rises of 2°C or less (Darrow et al., 2019), consistent with this result and our results.

Finally, a critical question—to which we cannot yet provide a definitive answer—is whether the mechanisms underlying the neuromodulatory effects of ultrasound are the same in our experiments and in *in vivo* experiments using low frequencies. Two results that strongly argue against similar mechanisms are the much lower intensities that have been reported to cause neuromodulatory effects in some low-frequency, *in vivo* experiments in small animal models (Tufail et al., 2010) as compared with our results, and the apparent increase in efficacy of ultrasound neuromodulation at lower frequencies in *in vivo* experiments. Both thermal and radiation force effects are proportional to ultrasound intensity, and to the ultrasound attenuation coefficient, which in tissue is proportional to frequency raised to the power of ∼1.1 (Hand, 1998), so these effects would generally be smaller in *in vivo* experiments (especially small animal experiments) as compared with our experiments. Compounding this issue is the fact that opposite dependences on the ultrasound frequency have been observed in *in vivo* and *in vitro* experiments. In an *in vivo* mouse model of ultrasound neuromodulation, it was determined that the efficacy of neuromodulation decreased with increasing frequency over the range 0.25-2.9 MHz (King et al., 2013; Ye et al., 2016). This frequency dependence is the opposite of what would be expected for either a thermal or radiation force mechanism. In contrast, Menz et al. (2019) found that the efficacy of neuromodulation increases with frequency over the range 1.9-43 MHz in the retina *in vitro*, a thin neural tissue preparation similar to the one used in our experiments. However, they proposed a model to explain this discrepancy. Lower ultrasound frequencies generally result in a larger stimulated tissue volume, which could translate into a more effective stimulus for certain structures of circuit-level neural connectivity, despite a weaker effect of low-frequency ultrasound at the level of an individual cell. (The model was presented in the context of a radiation force mechanism, but the same principle could apply for a thermal mechanism). The idea that circuit-level mechanisms can amplify the response to ultrasound is supported by comparison of our results with the response to ultrasound in the retina at 43 MHz. In the retina, potentiation of action-potential firing by ultrasound at 43 MHz saturates at 10 W/cm^2^, as measured at the population level in an intact, active neural circuit. Although we have not performed a detailed investigation of the intensity dependence, we find that a much higher intensity, 50 W/cm^2^, produces relatively moderate effects on excitability in single cells in the absence of significant network activity. Focusing solely on the local, cell-level amplitude of thermal and radiation force effects may therefore overlook important factors related to the global, network-level distribution of these effects. Such considerations may eliminate the differences in effective ultrasound parameters for *in vivo* and *in vitro* experiments as an argument against similar physical mechanisms for neuromodulation by high- and low-frequency ultrasound.

In conclusion, our results demonstrate that high-frequency ultrasound is a viable and promising modality for neuromodulation applications where frequency is not limited by transmission through the skull, and our insights into the common molecular mechanisms underlying both inhibitory and excitatory effects of high-frequency ultrasound pave the way for rational design and optimization of neuromodulation protocols to consistently produce either inhibitory or excitatory effects.

## ACKNOWLEDGMENTS

We thank the PIs and members of the Baccus, Butts-Pauly and Huguenard laboratories for helpful discussions. We thank Mike Menz and Stephen Baccus for critically reading the manuscript. We thank Marianna Kiraly, Dong Li, and Jason Clark for help with hippocampal slice preparation and recording. We thank the Stephen Boxer lab for sharing their plasma cleaner.

This work was supported by NIH R01 EB019005 (M.M. and B.K.Y.) and NIH BRAIN initiative R01 NS11215 (M.M. and B.K.Y.).

The authors declare no competing financial interests.

## AUTHOR CONTRIBUTIONS

Martin Prieto performed experiments, analyzed data, and wrote the manuscript. Martin Prieto and Kamyar Firouzi performed computational modeling. Merritt Maduke, Daniel Madison, and Butrus Khuri-Yakub supervised research. All authors contributed to the design and interpretation of experiments and edited and revised the manuscript.

## SUPPLEMENTAL MATERIALS

**Supplemental Figure 1.**
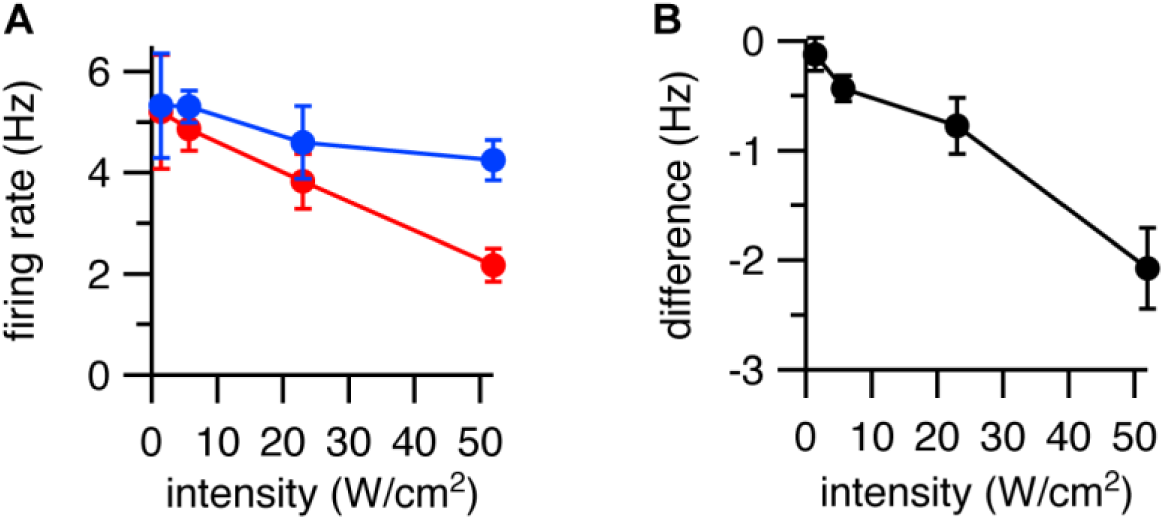
Effects of ultrasound at different intensities on firing rate. **A.** Mean (± SEM) action potential firing rate with (*red*) or without (*blue*) a 1-s ultrasound pulse at various intensities starting 200 ms after the start of a 2-s current step, as measured during the period of overlap between the ultrasound and current stimuli, or during the same time period in the absence of ultrasound. N = 4 cells, except for 6 W/cm^2^, where N = 3 cells. **B.** As in panel A, but showing the difference in firing rate between the ultrasound and control conditions.

**Supplemental Figure 2.**
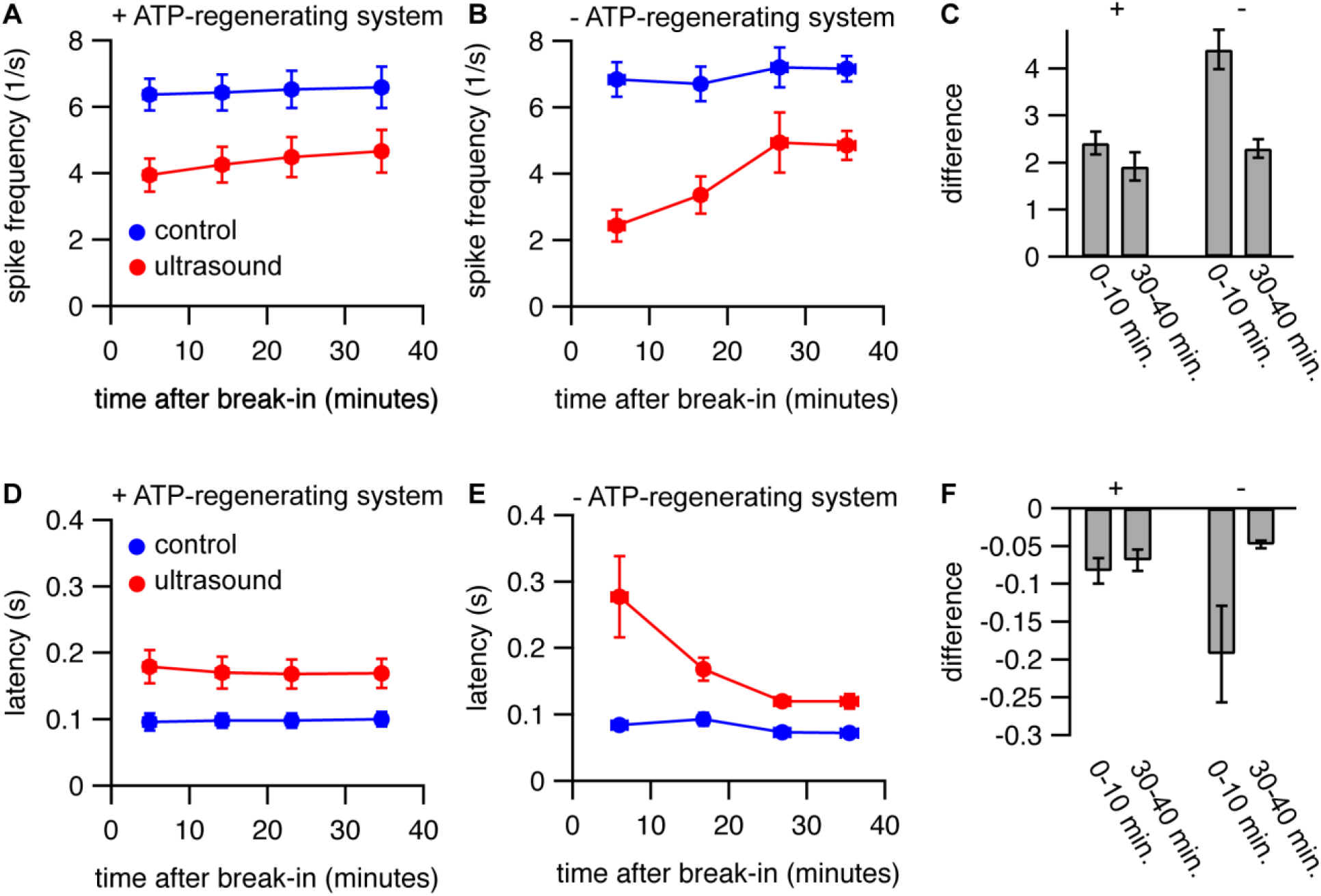
An ATP-regenerating internal solution stabilizes the response to ultrasound. **A-B.** Mean (±SEM) spike rates during the first 500 ms of a current step in the presence (*red circles*) and absence (*blue circles*) of a 1-s ultrasound application starting 500 ms before the current step, as a function of time relative to break-in (establishment of whole-cell recording configuration). The internal solution contained 10 Na-phosphocreatine with (A) or without (B) 50 U/mL creatine phosphokinase to provide an ATP-regenerating system. Spike rates were measured at various time points between 0 and 10, 10 and 20, 20 and 30, and 30 and 40 minutes after break-in. The x-values represent the mean start time for the protocol to measure spike rates (which comprised 2 minutes of recording time) with horizontal error bars (in some cases smaller than the symbol size) representing the SEM. The amplitude of the current step was adjusted over time to maintain spiking behavior as close as possible to that at the start of the experiment. **C.** Mean (±SEM) difference in spike rate between the control and ultrasound conditions for measurements at 1-10 minutes after break-in and 30-40 minutes after break-in, with (*left*) or without (*right*) the ATP-regenerating system. The difference was only statistically significant without the ATP-regenerating system (P = 0.11, with; and P = 0.0064, without). **D-F.** Same as A-B, but for latency to the first action potential following the start of the current step. N = 10 cells with and N = 6 cells without the ATP-regenerating system. The difference was not statistically significant for either group (P = 0.21, with; and P = 0.089, without).

**Supplemental Figure 3.**
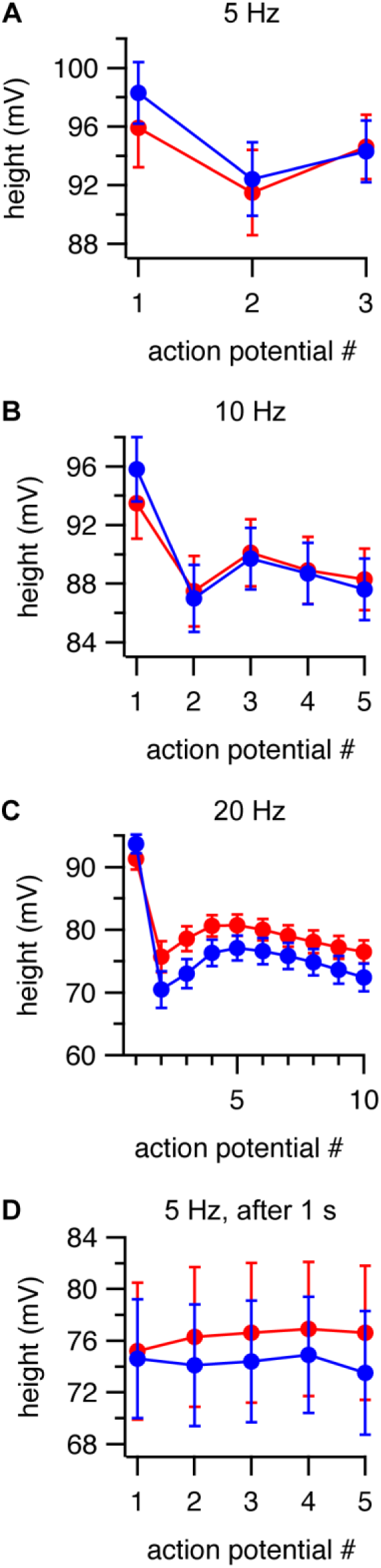
Effects of ultrasound on action potential height. **A-C.** Mean (±SEM) action potential heights as a function of action potential number in the presence (*red*) and absence (*blue*) of a 1-s ultrasound pulse starting 500 ms before the current step, for cells firing at an average firing rates (as measured during the first 500 ms of the current step) of approximately 5 Hz (N = 13), 10 Hz (N = 15), and 20 Hz (N = 13) in the control condition. **D.** As in A-C, but with ultrasound applied 1 s after the start of a 3-s current step, for cells firing at an average firing rate of approximately 5 Hz in the control condition, with firing rate determined in a 1-s window starting 1 s after the current step (corresponding to the time period of the ultrasound stimulus), and action potential number relative to the start of the ultrasound stimulus (N = 6). Note that the y-axes do not begin at zero.

**Supplemental Figure 4.**
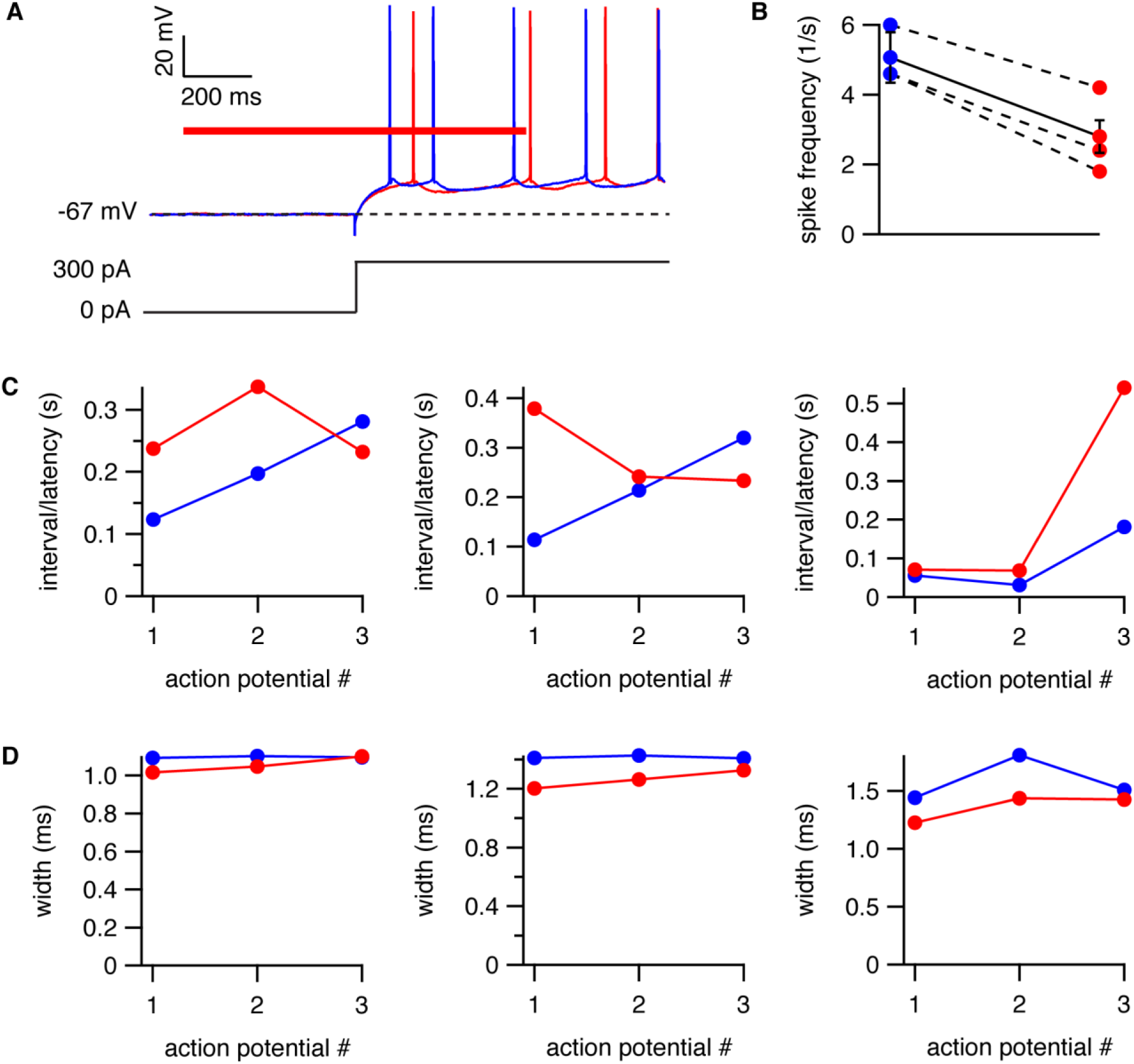
Effects of ultrasound on action potential firing and waveform at near-physiological temperature. **A.** Example voltage traces showing inhibition of action potential firing by ultrasound at 30°C. The response to a 300-pA current step is shown with (*red voltage trace*) and without (*blue voltage trace*) a 1-s ultrasound pulse (*red bar*) applied 500 ms before the start of the current step. The *dashed line* indicates the resting membrane voltage. **B.** Mean (± SEM, N =3) spike frequency during the first 500 ms of the current step (corresponding to the period of overlap between the current and ultrasound stimuli, or the equivalent time period in the absence of ultrasound) for the protocol shown in panel A for the control (*blue*) and ultrasound (*red*) conditions, for cells firing at an average spike frequency of approximately 5 Hz in the absence of ultrasound. Data points for the individual cells are shown connected by *dashed lines*. (Compare Figure 1E.) **C.** Latency between the start of the current step and the first action potential, and between the first and second, and second and third action potentials, for the control (*blue*) and ultrasound (*red*) conditions for three individual cells. (Compare Figure 4A.) **D.** As in panel C, but for effects of ultrasound on action potential width. (Compare Figure 6A.)

